# Domain adaptive neural networks improve cross-species prediction of transcription factor binding

**DOI:** 10.1101/2021.02.13.431115

**Authors:** Kelly Cochran, Divyanshi Srivastava, Avanti Shrikumar, Akshay Balsubramani, Ross C. Hardison, Anshul Kundaje, Shaun Mahony

## Abstract

The intrinsic DNA sequence preferences and cell-type specific cooperative partners of transcription factors (TFs) are typically highly conserved. Hence, despite the rapid evolutionary turnover of individual TF binding sites, predictive sequence models of cell-type specific genomic occupancy of a TF in one species should generalize to closely matched cell types in a related species. To assess the viability of cross-species TF binding prediction, we train neural networks to discriminate ChIP-seq peak locations from genomic background and evaluate their performance within and across species. Cross-species predictive performance is consistently worse than within-species performance, which we show is caused in part by species-specific repeats. To account for this domain shift, we use an augmented network architecture to automatically discourage learning of training species-specific sequence features. This domain adaptation approach corrects for prediction errors on species-specific repeats and improves overall cross-species model performance. Our results demonstrate that cross-species TF binding prediction is feasible when models account for domain shifts driven by species-specific repeats.

---

Characterizing where transcription factors (TFs) bind to the genome, and which genes they regulate, is key to understanding the regulatory networks that establish and maintain cell identity. A TF’s genomic occupancy depends not only on its intrinsic DNA sequence preferences, but also on several cell-specific factors, including local TF concentration, chromatin state, and cooperative binding schemes with other regulators (Siggers and Gordân 2014; Slattery et al. 2014; Srivastava and Mahony 2020). Experimental assays such as ChIPseq can profile a TF’s genome-wide occupancy within a given cell type, but such experiments remain costly, rely on relatively large numbers of cells, and require either high-quality TF-specific antibodies or epitope tagging strategies (Park 2009; Savic et al. 2015). Accurate predictive models of TF binding could circumvent the need to perform costly experiments across all cell types and all species of interest.

Cross-species TF binding prediction is complicated by the rapid evolutionary turnover of individual TF binding sites across mammalian genomes, even within cell types that have conserved phenotypes. For example, only 12-14% of binding sites for the key liver regulators CEBPA and HNF4A are shared across orthologous genomic locations in mouse and human livers (Schmidt et al. 2010). On the other hand, the general features of tissue-specific regulatory networks appear to be strongly conserved across mammalian species. The amino acid sequences of TF proteins, their DNA binding domains, and intrinsic DNA sequence preferences are typically highly conserved (e.g., both CEBPA and HNF4A have at least 93% whole protein sequence identity between human and mouse). Further, the same cohorts of orthologous TFs appear to drive regulatory activities in homologous tissues. Thus, while genome sequence conservation information is not sufficient to accurately predict TF binding sites across species, it may still be possible to develop predictive models that learn the sequence determinants of cell-type specific TF binding and generalize across species. Indeed, several recent studies have demonstrated the feasibility of cross-species prediction of regulatory profiles using machine learning approaches (Chen et al. 2018; Kelley 2020; Schreiber et al. 2020; Huh et al. 2018).

Here, we evaluate different training strategies on the generalizability of neural network models of cell-type specific TF occupancy across species. We train our model using genome-wide TF ChIP-seq data in a given cell type in one species, and then assess its performance in predicting genome-wide binding of the same TF in a closely matched cell type in a different species. Specifically, we focus on predicting binding of four TFs (CTCF, CEBPA, HNF4A, and RXRA) in liver due to the existence of high quality ChIP-seq data in both mouse and human. The models for all TFs showed higher predictive performance for training and test sets from the same species as compared to training and test sets from different species. We show that one source of this cross-species performance gap is a systematic misclassification of transposable elements that are specific to the target species (and which were thus unseen during model training).

We further demonstrate that integrating an unsupervised domain adaptation approach into model training partially addresses the cross-species performance gap. Our domain adaptation strategy involves a neural network architecture with two sub-networks that share an underlying convolutional layer. We train the two sub-networks in parallel on different tasks. One subnetwork is trained with standard backpropagation to optimize classification of TF bound and unbound sequences in one species (the source domain). The other subnetwork attempts to predict species labels from sequences drawn randomly from two species (the source and target domain), but training is subject to a gradient reversal layer (GRL) (Ganin et al. 2016). While backpropagation typically has the effect of giving higher weights to discriminative features, a GRL reverses this effect, and discriminative features are down-weighted. Thus, our network encourages features in the shared convolutional layer that discriminate between bound and unbound sites, while simultaneously discouraging features that are species-specific. Importantly, we neither need nor use TF binding labels from the target species at any stage in training. We show that domain adaptation techniques have the potential to improve cross-species TF binding prediction, particularly by preventing misprediction on species-specific repeats.

## Results

### Conventionally trained neural network models of TF binding show reduced predictive performance across species

First, we set out to evaluate the ability of neural networks to predict TF binding in a previously unseen species. We chose neural networks due to their ability to learn arbitrarily complex predictive sequence patterns (Avsec et al. 2021a; Avsec et al. 2021b; Fudenberg et al. 2020; Kelley 2018; Koo et al. 2021). In particular, hybrid convolutional and recurrent network architectures have successfully been applied to accurately predict TF binding in diverse applications (Quang and Xie 2016; Quang and Xie 2019; Srivastava et al. 2020). The motivation behind these architectures is that convolutional filters can encode binding site motifs and other contiguous sequence features, while the recurrent layers can model flexible, higher-order spatial organization of these features. Our baseline neural network is designed in line with these state-of-the-art hybrid architectures (Figure 1).

**Figure 1:**
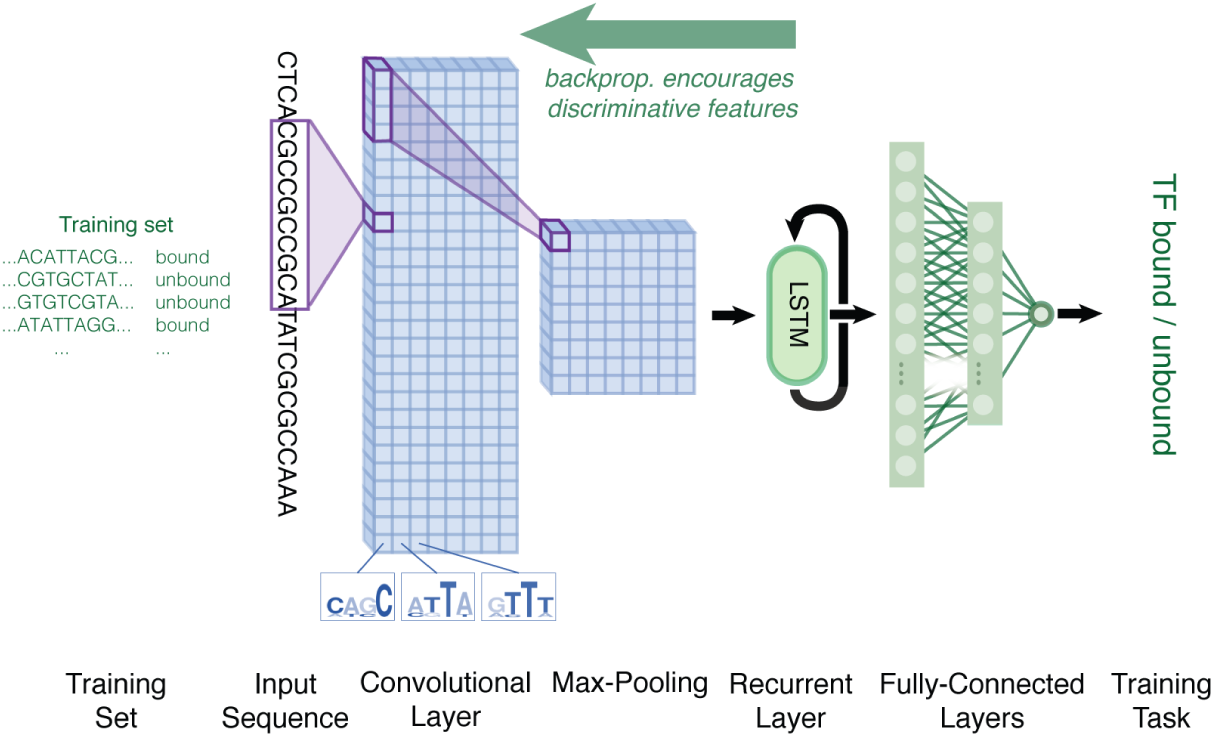
Conventional network architecture. Convolutional filters scan the 500-bp input DNA sequence for TF binding features. The convolutional layer is followed by a recurrent layer (LSTM) and two fully connected layers. A final sigmoid-activated neuron predicts if a ChIP-seq peak falls within the input window.

Using this architecture, named the “conventional model,” we trained the network to predict whether a given input sequence contained a ChIP-seq peak or not, using training data from a single source species, and then assessed the model’s predictive performance on entire held-out chromosomes in both the source species and a target (previously unseen) species. We chose mouse and human as our species of interest due to the availability of high-quality TF ChIP-seq datasets in liver from both species and the high conservation of key regulator TFs present in both species. For four different TFs, we trained two sets of models: one with mouse as the source species, and the other with human as the source species. To monitor reproducibility, model training was repeated 5 times for each TF and source species.

As models trained for 15 epochs, we monitored source-species and target-species performance on heldout validation sets (Figure 2). Performance was measured using the area under the precision-recall curve (auPRC) which is sensitive to the extreme class imbalance of labels in our TF binding prediction task. We observed that over the course of model training, improvements in source-species auPRC did not always translate to improved auPRC in the target species. Overall, cross-species auPRC showed greater variability across epochs and model replicates compared to source-species auPRC. For two TFs, CEBPA and HNF4A, the mouse-trained models’ performance on the human validation set appeared to split part way through training – based on cross-species auPRC, some model-replicates appeared to become trapped in a suboptimal state relative to other models (see divergence in orange lines in left column of Figure 2); meanwhile, the training-species auPRC did not show a similar trend. Evidently, validation set performance in the source species is not a reliable surrogate for validation set performance in the target species.

**Figure 2:**
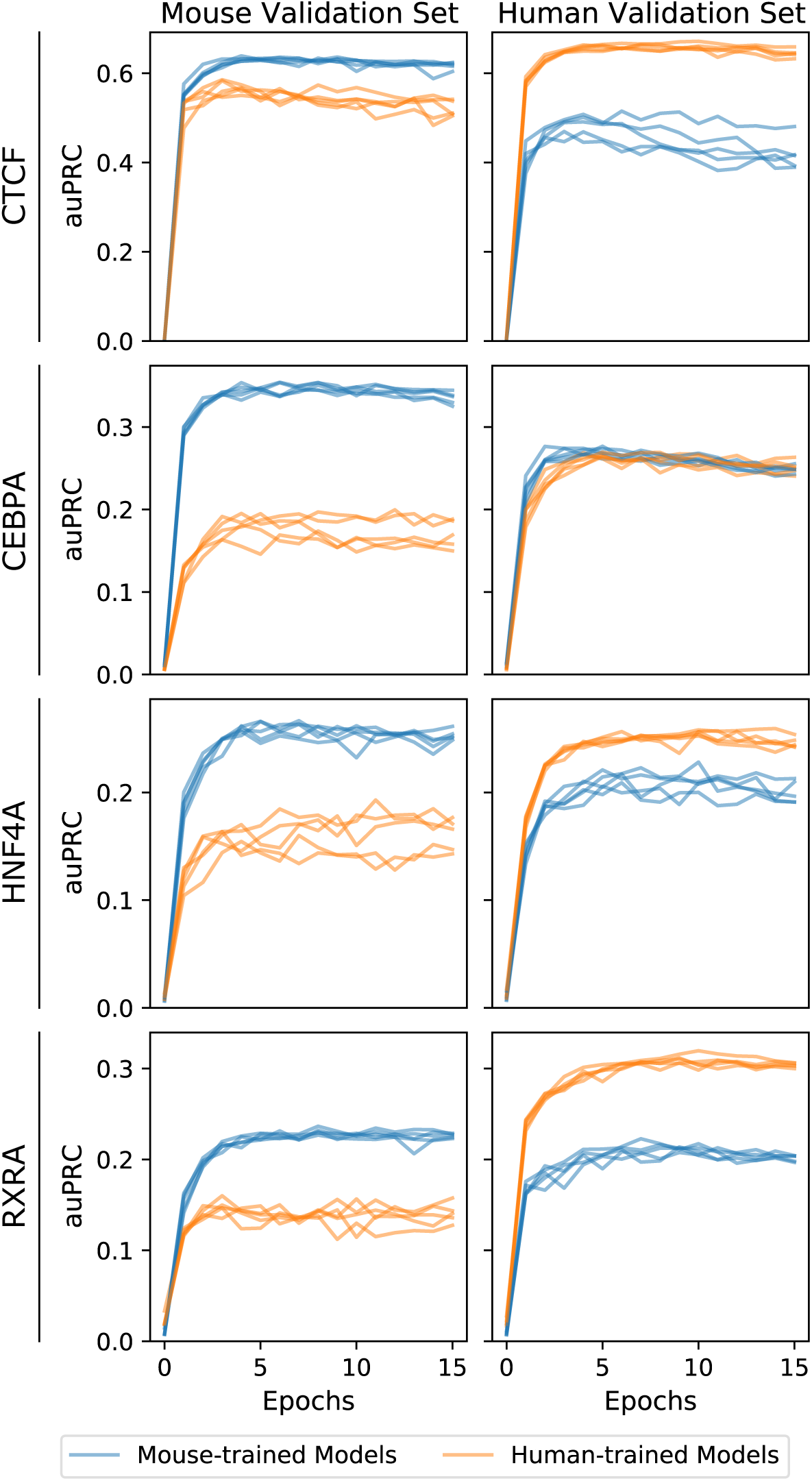
Model performance over the course of training, evaluated on held-out validation data from mouse (left) and human (right) Chromosome 1. Five models were independently trained for each TF and source species (mouse-trained models in blue, human-trained models in orange). Values at epoch 0 are evaluations of models after weight initialization but before training (akin to a random baseline). Note that auPRCs are not directly comparable between different validation sets because ground truth labels are derived from a different experiment for each dataset; the auPRC will depend on the fraction of sites labeled bound as well as model prediction correctness.

Nevertheless, the epochs where models had highest source-species auPRCs were often epochs where models had near-best cross-species auPRC. Thus, we selected models saved at the point in training when source-species auPRC was maximized for downstream analysis. We next evaluated performance on held-out test datasets (distinct from the validation datasets) from each species (Figure 3).

**Figure 3:**
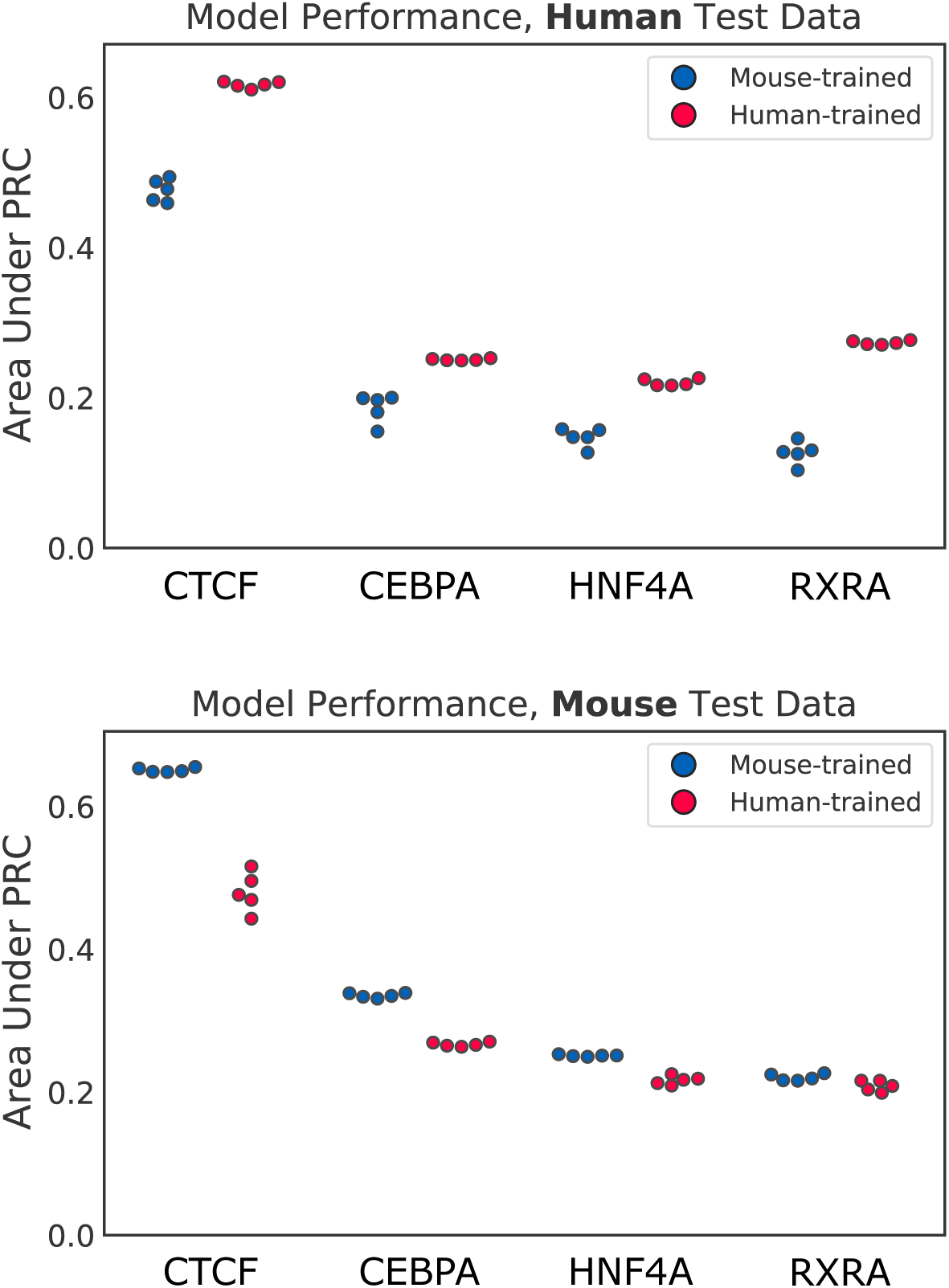
Model performance evaluated on held-out test data: Chromosome 2 from human (top) and mouse (bottom). Five models were independently trained for each TF and source species.

We observe across all TFs that for a given target species, the models trained in that species always outperformed or matched the performance of the models trained in the other species. We refer to this within-species vs. cross-species auPRC difference as a crossspecies performance gap, while noting that models trained in either species were still relatively effective at cross-species prediction. Intriguingly, the cross-species gap was wider for mouse-trained models predicting in human than for human-trained models predicting in mouse. For this reason, subsequent analysis focuses on addressing the mouse-to-human gap.

To get a sense of how specific to our model design or training strategy this cross-species gap might be, we sought to apply a sufficiently different machine learning approach to the same problem and datasets and assess whether the cross-species gap persists. We trained gapped *k*-mer support vector machines, or gkSVMs, to classify a balanced sample of bound vs. unbound windows for each TF and species (Ghandi et al. 2014; Lee 2016). We then evaluated those models on the set of nonoverlapping windows in each test dataset (Figure S1). The gkmSVM auPRC values are drastically lower than those of the neural networks across the board, demonstrating that our deep learning approach can indeed out-perform related methods on this task. We also observe that the cross-species gap persists, although it shrinks in absolute magnitude, presumably due to the much lower auPRC values overall.

### The mouse-to-human cross-species gap originates from misprediction of both bound and unbound sites

Since the target-species model consistently outperforms the source-species model (on target-species validation), there must be some set of differentially predicted sites that the target-species model predicts correctly, but the source-species model does not. By comparing the distribution of source-model and target-model predictions over all target-species genomic windows, we can potentially identify trends of systematic errors unique to the source-species model. Whether these differentially predicted sites are primarily false positives (unbound sites incorrectly predicted to be bound), false negatives (bound sites incorrectly predicted as unbound), or a combination of both can provide useful insight into the performance gap between the source and target models.

For each TF, we generated predictions over the genomic windows in the human test dataset from both our mouse-trained and human-trained models. Then, we plotted all of the human-genome test sites using the average mouse model prediction (over 5 independent training runs) and the average human model prediction as the xand y-axis, respectively (Figure 4). Bound and unbound sites are segregated into separate plots for clarity.

**Figure 4:**
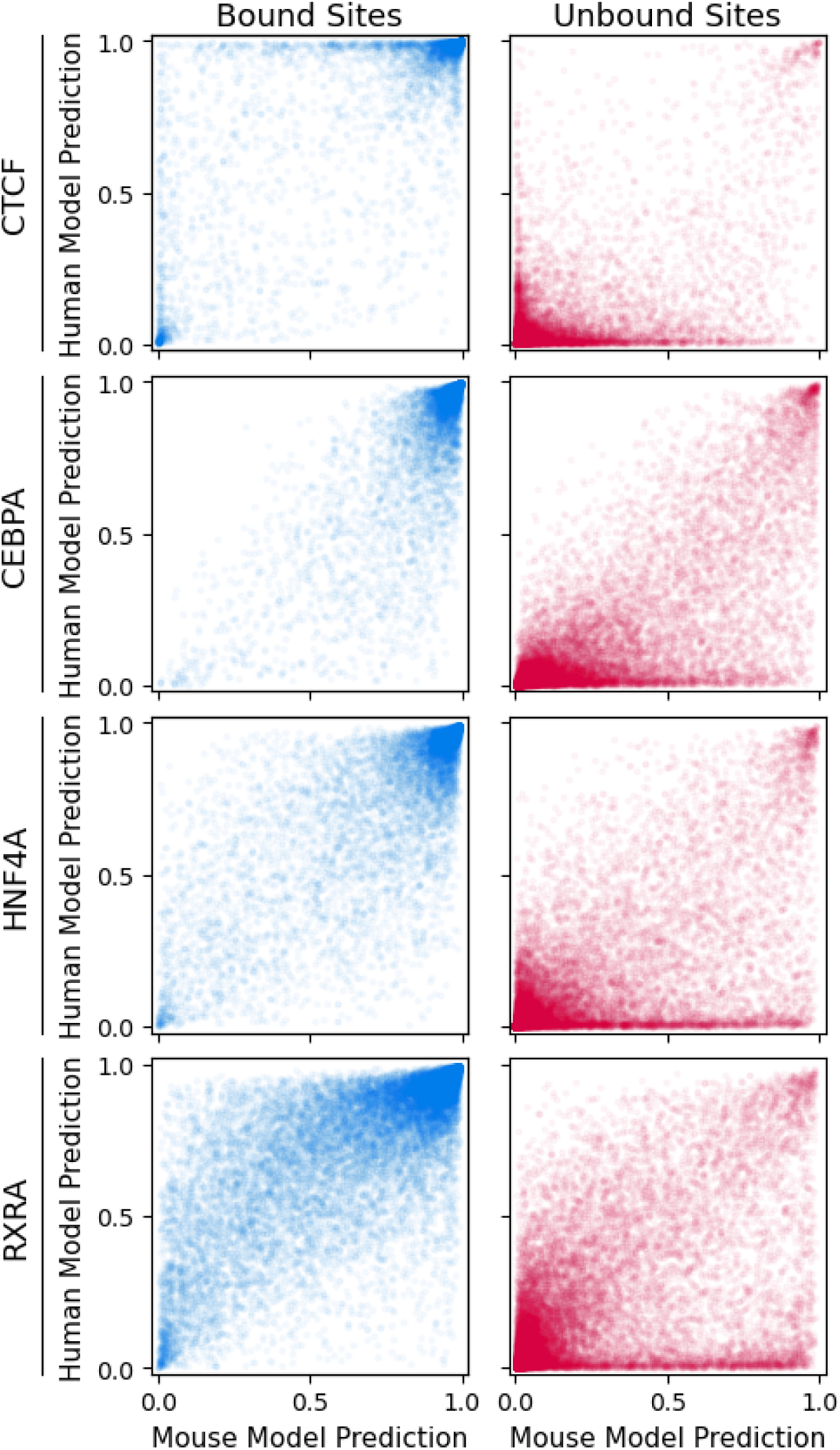
Both bound and unbound sites from human Chromosome 2 show evidence of differential binding predictions by human-trained (y-axis) vs. mouse-trained (x-axis) models. For visual clarity, only 25% of bound sites and 5% of unbound sites are shown (sampled systematically).

For all TFs, the unbound site plots show a large set of windows given low scores by the human model but mid-range to high scores by the mouse model – these are false positives unique to cross-species prediction (Figure 4 right column, bottom/bottom-right region of each plot). These sites are distinct from false positives mistakenly predicted highly by both models, as those common false positives would not contribute significantly to the auPRC gap. Additionally, in the bound site plots of all TFs except CEBPA, we see some bound sites that are scored high by the human model but are given midrange to low scores by the mouse model – these are crossspecies-unique false negatives (Figure 4 left column, top left region of each plot). Hence, our cross-species models are committing prediction errors in both directions on separate sets of sites. The errors for the unbound sites appear more prevalent than the errors for the bound sites.

### Motif-like sequence features discriminate between true-positive and false-negative mouse model predictions

Since the only input to our models is DNA sequence, sequence features must be responsible for differential prediction of certain sites across source and target models. Other potential culprits, such as chromatin accessibility changes or co-factor binding, may contribute to TF binding divergence across species without changes to sequence; but without an association between those factors and sequence, the human-trained model would not be able to gain an advantage over the mouse-trained model by training on sequence input alone. Thus, we focused on genomic sequence to understand differential site prediction.

To begin, we searched for sequence features associated with differential prediction of bound sites from the human genome – specifically, we compared bound sequences that both the human-trained and mouse-trained models correctly predicted (true positives) to bound sequences the human-trained model correctly predicted but the mouse-trained model did not (mouse-specific false negatives). We used SeqUnwinder, a tool for deconvolving discriminative sequence features between sets of genomic sequences, to extract motifs that can discriminate between the two groups of sequences and quantitatively assess how distinguishable the sequence groups are (Kakumanu et al. 2017). SeqUnwinder was able to distinguish mouse-specific false negatives from true positives and randomly selected background genomic sequences with area under the ROC curve (au-ROC) of 0.84, 0.74, 0.83, and 0.88 for CTCF, CEBPA, HNF4A, and RXRA, respectively. Figure S2 shows the breakdown of sequence features that are able to distinguish between mouse-specific false negatives and true positives for each TF. Thus, we were able to identify TF-specific motifs that were enriched (or depleted) at mouse-specific false negatives. However, we did not observe systemic sequence features that unanimously contributed to the performance gap across all TFs studied, beyond a poly-A/poly-T motif.

### Primate-unique SINEs are a dominant source of the mouse-to-human cross-species gap

One potential source of sequences that could confuse a cross-species model are repeat elements found in the genome of the target species but not the source species. *Alu* elements, a type of SINE, cover a large portion (10%) of the human genome and are found only in primates (Batzer and Deininger 2002). Several other factors make *Alus* even more likely candidates for confounding mouse-to-human TF binding predictions: they are enriched in gene-rich, GC-rich areas of the genome and contain 33% of the genome’s CpG dinucleotides (a marker for promoter regions); they may play a role in gene regulation; and in silico studies have previously found putative TF binding sites within *Alu* sequences (Batzer and Deininger 2002; Schmid 1998; Ferrari et al. 2019; Polak and Domany 2006).

Figure 5 shows only the unbound human-genome windows that overlap annotated *Alu* elements. Table 1 provides corresponding quantification of *Alu* enrichment. Note that while *Alu* elements are typically poorly mappable, and it is thus often difficult to assign them as bound or unbound in ChIP-seq experiments, we focus analyses here only on highly mappable *Alu* instances (see Methods). Across all four TFs, we see that *Alus* are substantially enriched in the unbound windows predicted incorrectly only by the mouse model. On average, 83% of these false positives unique to the mouse model overlap with an *Alu* element, compared to the average overlap rate of 21% for unbound sites overall, or 17% for unbound sites incorrectly predicted by both models. In contrast, *Alus* on average only overlap 7% of false negatives unique to the mouse model, which is less than the overlap fraction for bound sites overall (14%) and for false negatives common to both models (10%). We repeated this analysis using other repeat classes, including LINEs and LTRs, and confirmed that no other major repeat family shows an enrichment of comparable strength with either the false positives or false negatives unique to the mouse model (Table S1). Investigating the enrichment of individual *Alu* subfamilies in mouse-model-unique false positives showed that this phenomenon is not restricted to a single subtype of *Alu*, but that subfamilies are enriched at different levels in a manner that is TF-specific and varies particularly between the *AluJ, AluS*, and *AluY* subfamily groupings (Figure S3).

**Table 1:** Percent of windows overlapping an *Alu* element, for various categories of genomic windows from the held-out test set. *Alu* elements dominate the false positives unique to the mouse models. FPs: false positives. FNs: false negatives. See Methods for more details on site categorization.

| TF | Bound | FN (Both Models) | FN (Mouse Only) | Unbound | FP (Both Models) | FP (Mouse Only) |
| --- | --- | --- | --- | --- | --- | --- |
| CTCF | 12.0% | 10.7% | 9.6% | 21.2% | 8.5% | **65.2%** |
| CEBPA | 18.3% | 11.5% | 0.0% | 21.2% | 23.2% | **82.2%** |
| HNF4A | 13.6% | 9.1% | 6.9% | 21.3% | 16.0% | **87.3%** |
| RXRA | 13.3% | 9.7% | 4.7% | 21.3% | 19.2% | **96.9%** |

**Figure 5:**
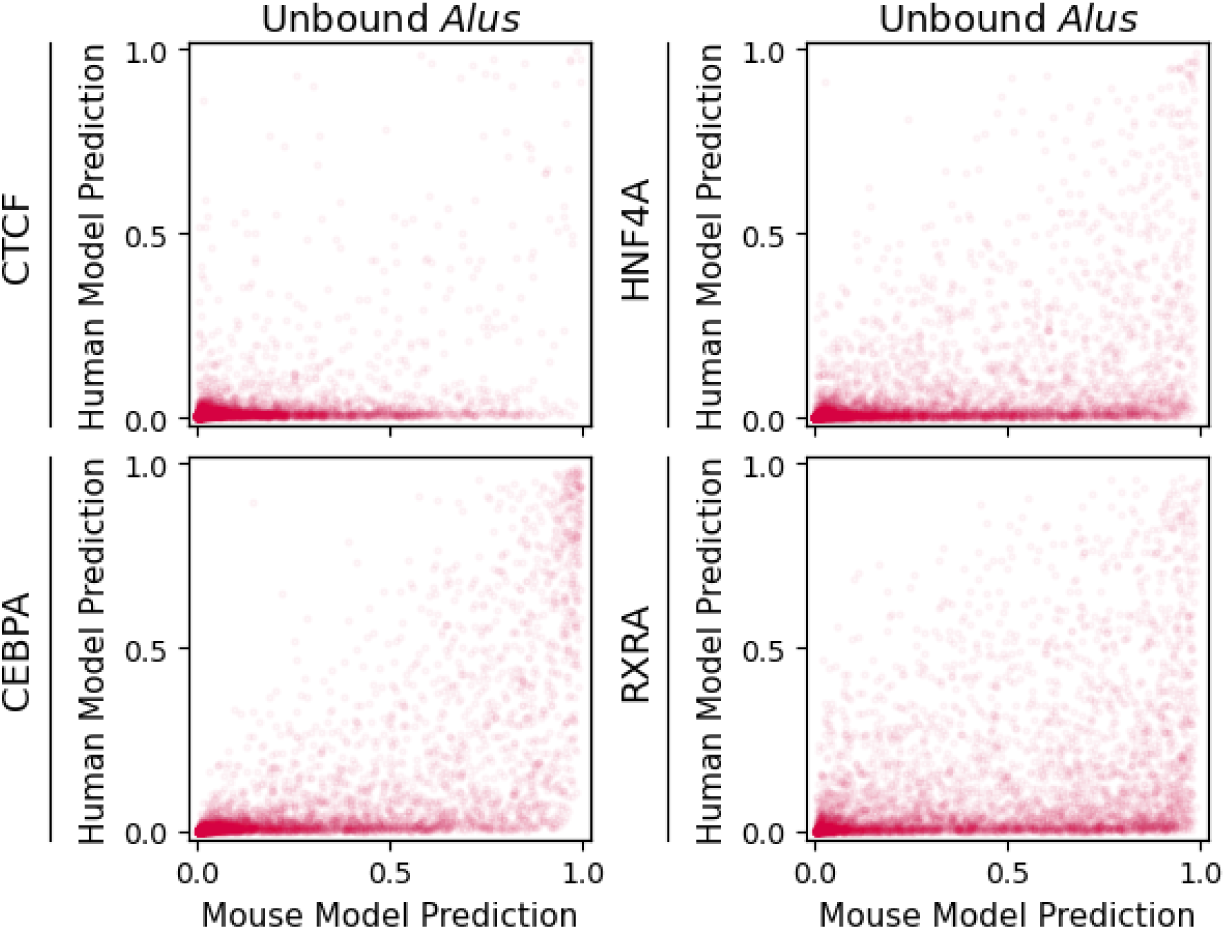
Most unbound sites from the human genome mispredicted by mouse-trained models (x-axis), but not by human-trained (y-axis) models, contain *Alu* repeats. For visual clarity, only 5% of windows are shown.

Thus, the vast majority of the false positives from the human genome mispredicted only by mouse models can be directly attributed to one type of primate-unique repeat element. We did not observe any similar direct associations between primate-unique elements and the false negatives unique to the mouse model, besides the expected depletion of *Alu* elements.

### Model interpretation reveals sequence features driving divergent mouse and human model predictions

To understand why mouse and human models make divergent predictions at some sites, we compared basepair resolution importance scores from both models at selected example sites. Specifically, we implemented a strategy similar to in silico mutagenesis (ISM) where a base’s score was determined by the differential model output between the original sequence and the sequence with 5bp centered on that base replaced with bases from a dinucleotide-shuffled reference (Alipanahi et al. 2015). We observed that this strategy outperformed backpropagation-based scoring methods, potentially by avoiding gradient instability.

First, we compared importance scores between the mouse and human models at example bound sites that both models predicted correctly (Figure S4). If the two models learned to use similar logic to make binding predictions, we would expect to see similar sequence features highlighted in the importance scores. Throughout the tested examples across all four TFs, we observe that the scores generated by the mouse and human models are remarkably concordant. In particular, instances of the primary cognate motifs for the appropriate TF are highlighted by both models.

Next, we repeated the analysis on example unbound windows classified as mouse-model-unique false positives (Figure S5). At these sites, the mouse model’s predictions overshoot those of the human model by at least 0.5. Importance scores in this set of sites show much greater disagreement between the two models. Commonly across all four TFs, we observed two trends: first, the mouse models often assigned high importance to motif-sized contiguous stretches of bases which were not similarly recognized by the human models. These pseudo-motifs can superficially resemble approximate matches to the TF’s cognate motif. Second, the human models commonly showed apparent sensitivity to specific, sparse features which received negative scores of moderate to high magnitude. These observations imply that the human model has learned to ignore certain sequence features that the mouse model’s scores suggest are favorable for binding. Furthermore, the human model may be adopting that strategy based on whether or not there are nearby sequence contexts that indicate that the sequence is not a binding site.

### Human models trained without SINE examples behave like hybrid mouse-human models

To further characterize how *Alu* elements are influencing cross-species model performance, we trained additional models on the human dataset after removing all windows from the training dataset that overlap with any SINEs (Figure 6). We filtered out all SINEs, including the primate-specific *FLAM* and *FRAM* repeats as well as *Alus*, to avoid keeping examples that shared any sequence homology with *Alus*. The no-SINE models were evaluated on the same held-out chromosome test data used previously (which includes SINEs). For all TFs except CTCF, the no-SINE models perform substantially worse than models trained using the complete human training sets.

**Figure 6:**
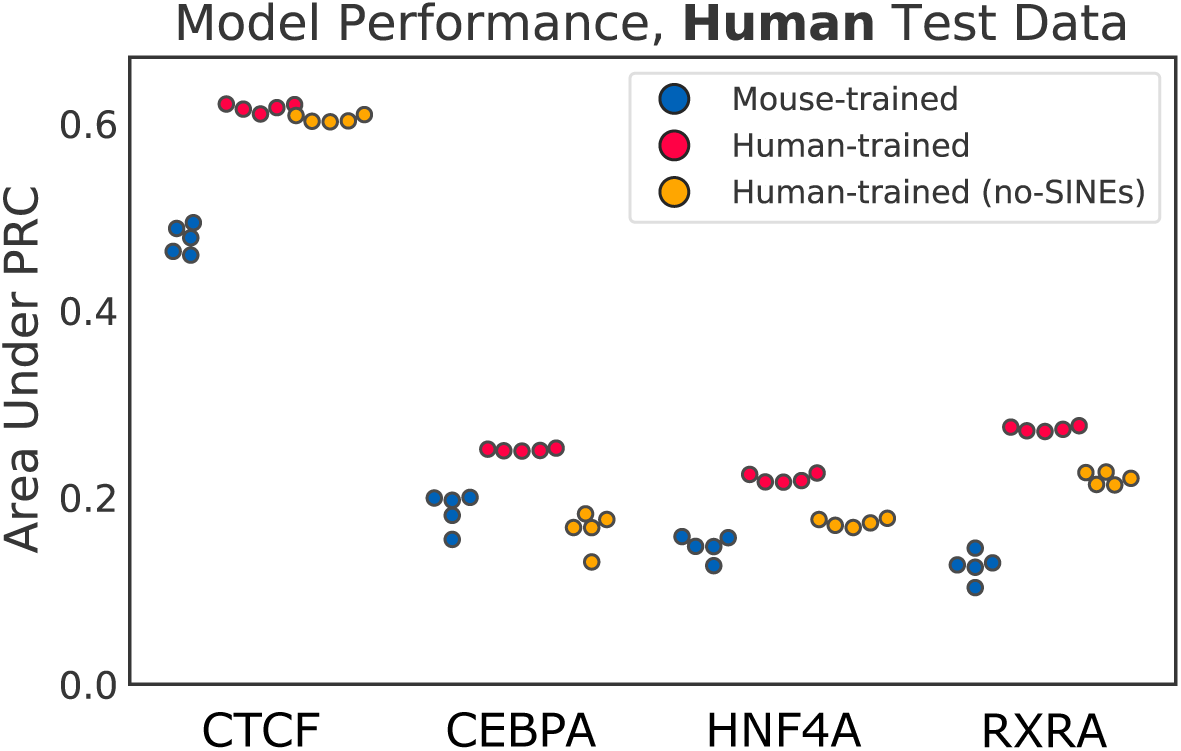
Performance of models that are mouse-trained (blue), human-trained with SINE examples (red), and human-trained without SINE examples (yellow), evaluated on the held-out human Chromosome 2. Five models were independently trained for each TF and training species.

Site-distribution plots show that, for unbound sites, no-SINE human-trained models make mispredictions in a pattern similar to mouse-trained models; there is a similarly-sized subset of unbound sites mispredicted by the no-SINE human-trained models but not by the standard human-trained models (Figure 7). Plotting only the sites that overlap with *Alus* confirms that the false positives unique to the no-SINEs model are predominantly *Alu* elements (Figure S6). For bound sites, on the other hand, no-SINE human-trained models make predictions that generally agree with predictions from standard human-trained models.

**Figure 7:**
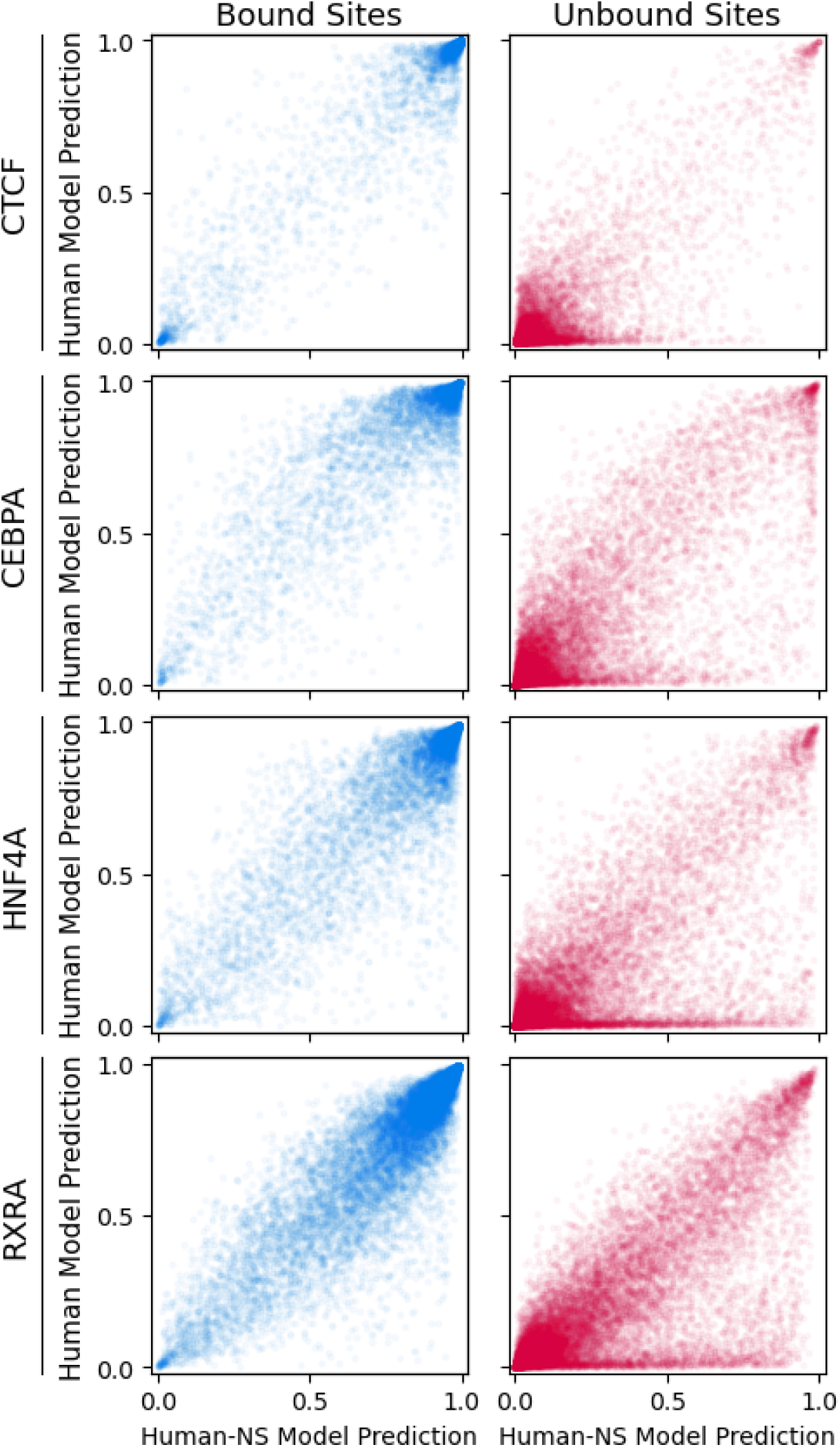
Differential human Chromosome 2 site predictions between models trained on human data with or without any examples of SINE windows. Human-NS: models trained on human data with no SINE examples. Similar to mouse-trained models, no-SINE human-trained models systematically mispredict some unbound sites.

This suggests that the *Alu* false positives unique to the mouse-trained model may simply be due to the fact that mouse models are not exposed to *Alus* during training (i.e., *Alu* elements are “out of distribution”). In addition, the reduction in model-unique false negatives observed when the no-SINE human-trained model is compared to the normal human-trained model suggests that those mispredictions are unrelated to *Alus*.

### Domain-adaptive mouse models can improve crossspecies performance

Having observed an apparent “domain shift” across species, partially attributable to species-unique repeats, our next step is to ask how we might bridge this gap and reduce the difference in cross-species model performance. Our problem is analogous to one encountered in some image classification tasks, where the test data is differently distributed from the training data to the extent that the model performs well on training data but much worse on test data (for example, the training images were taken during the day but the test images were taken at sunset). In these situations, various techniques for explicitly forcing the model to adapt across different image “domains” have been shown to improve performance at test time (e.g., Long et al. 2015; Bousmalis et al. 2016; Sun et al. 2016).

One unsupervised domain adaptation method utilizes a gradient reversal layer to encourage the “feature generator” portion of a neural network to be domain-generic (Ganin et al. 2016). The gradient reversal layer’s effect is to backpropagate a loss to the feature generator that prevents any domain-unique features from being learned. We chose to test the effectiveness of this version of domain adaptation for our cross-species TF binding prediction problem because we have observed evidence that domain-unique features (species-unique repeat elements) were a major component of the cross-species domain shift.

We modified our existing model architecture to perform training-integrated domain adaptation across species (Figure 8). A gradient reversal layer (GRL) was added in parallel with the LSTM, taking in the result of the max-pooling step (after the convolutional layer) as input. During standard feed-forward prediction, the GRL merely computes the identity of its input, but as the loss gradient backpropagates through the GRL, it is reversed. The output of the GRL then passes through two fully connected layers before reaching a new, secondary output neuron. This secondary output, a “species discriminator,” is tasked with predicting whether the model’s input genomic window is from the source or target species. The model training process is modified so that the model is exposed to sequences from both species, but only the binding labels of the source species (see Methods). Without the GRL, adding the species discrimination task to the model would encourage the convolutional filters to learn sequence features that best differentiate between the two species – features like species-unique repeats – but with the GRL included, the convolutional filters are instead *discouraged* from learning these features. We hypothesize that this domain-adaptive model will outperform our basic model architecture by reducing mispredictions on species-unique repeats.

**Figure 8:**
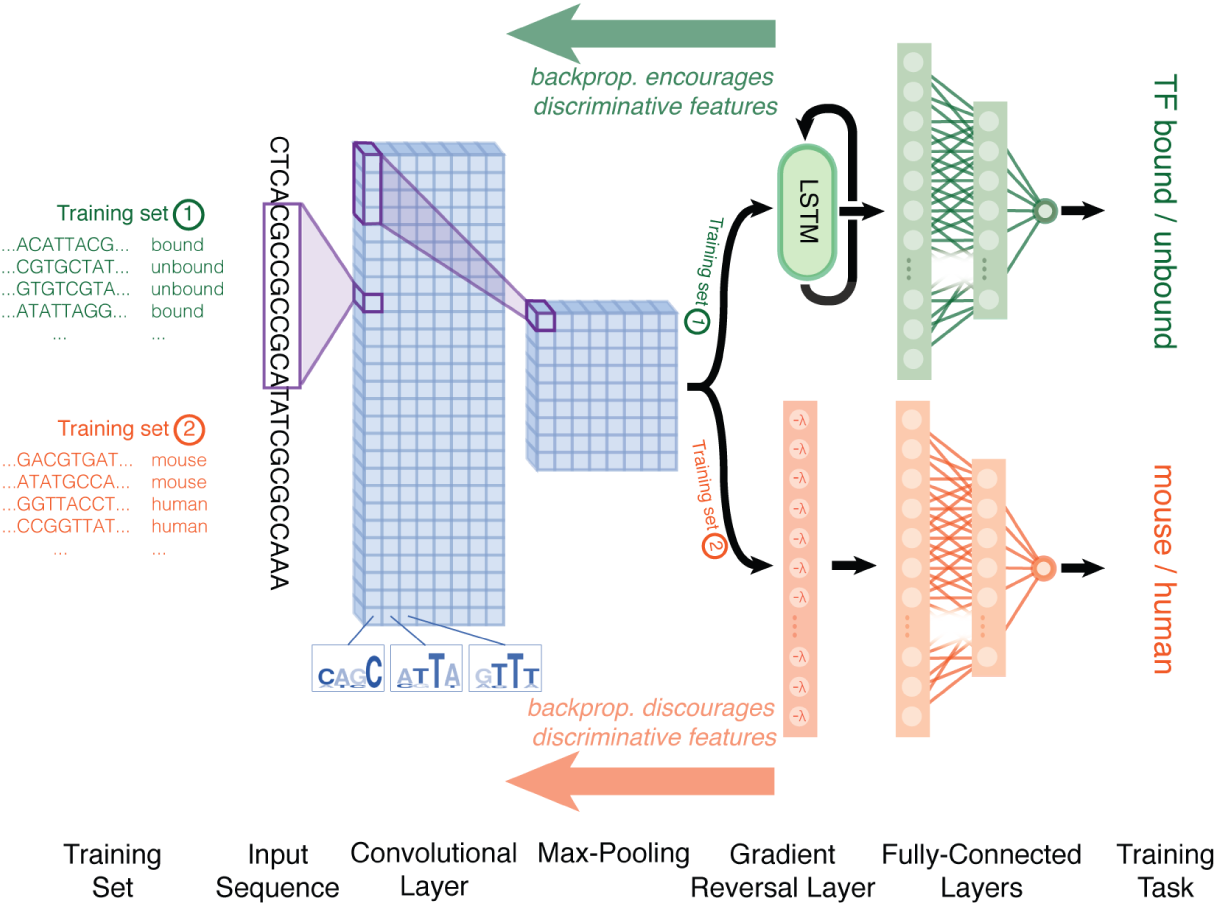
Domain-adaptive network architecture. The top network output predicts TF binding, as before, while the bottom network output predicts the species of origin of the input sequence window. The gradient reversal layer has the effect of discouraging the convolutional filters before it from learning sequence features relevant to the species prediction task.

We trained domain-adaptive models using the same binding training datasets as before and evaluated performance with the same held-out datasets. We observe that the auPRC for our domain-adaptive models on cross-species test data is moderately higher than the auPRC for the basic mouse models, for all TFs except CTCF, where auPRCs are merely equal (Figure 9, top, blue vs. green boxplots). The domain-adaptive models’ auPRCs on mouse test data, meanwhile, is comparable to the auPRCs of basic models (Figure 9, bottom, blue vs. green). While the auPRC improvement is promising, it is also modest in comparison to the full cross-species gap; the domain-adaptive models still do not achieve a level of performance comparable to same-species models (Figure 9, top, green vs. red).

**Figure 9:**
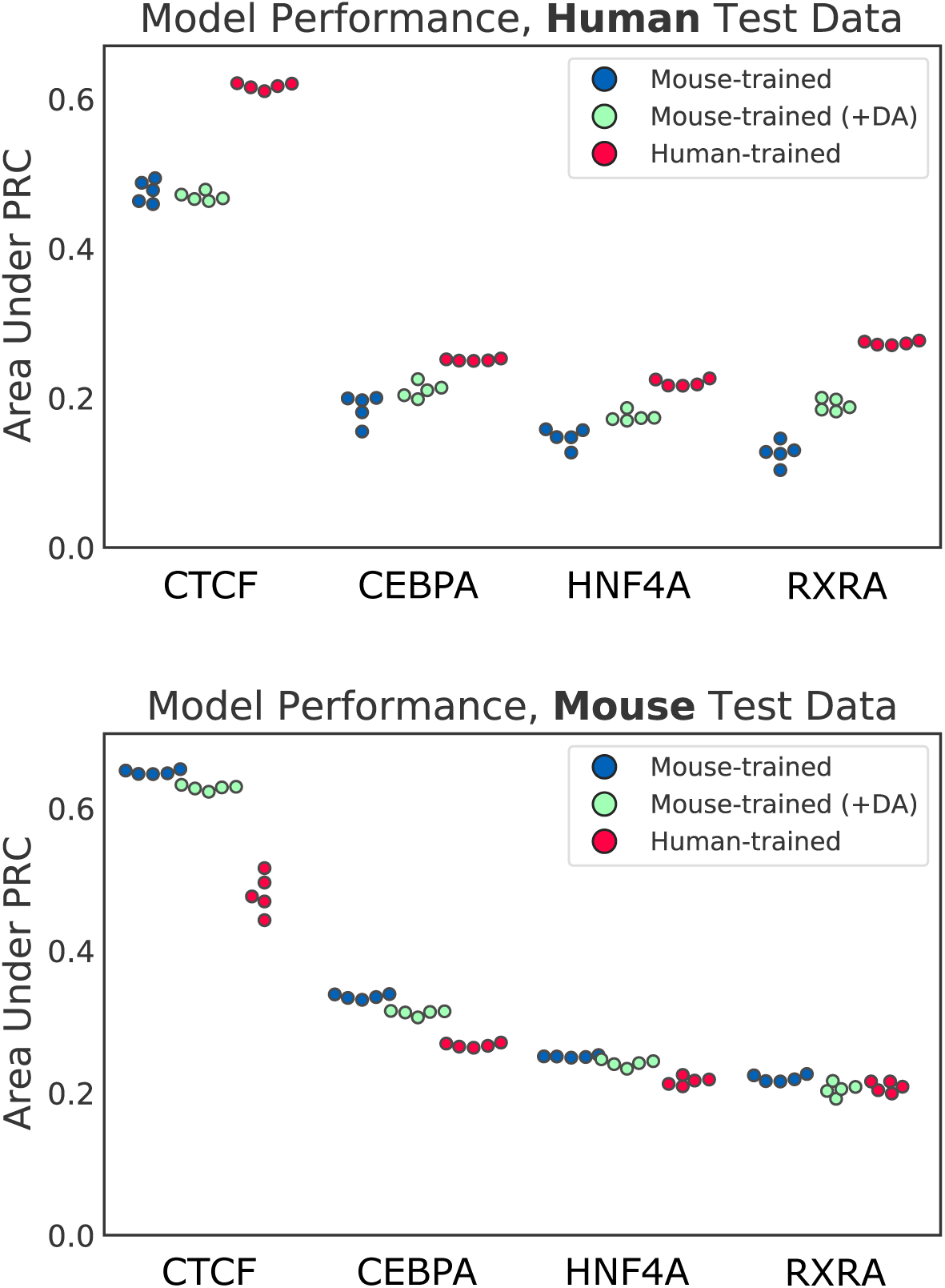
Performance of mouse-trained generic (blue), mouse-trained domain-adaptive (green), and human-trained (red) models, evaluated on human (top) and mouse (bottom) Chromosome 2. Five models were independently trained and evaluated for each TF and training species.

### Domain-adaptive mouse models reduce overprediction on *Alu* elements

Next, we repeated our site-distribution analysis to determine what constituted the domain-adaptive models’ improved performance. The unbound site plots in Figure 10 compare human genome predictions between domain-adaptive mouse models and the original human models. *Alu* elements are highlighted in Figure 11, with quantification in Table S2.

**Figure 10:**
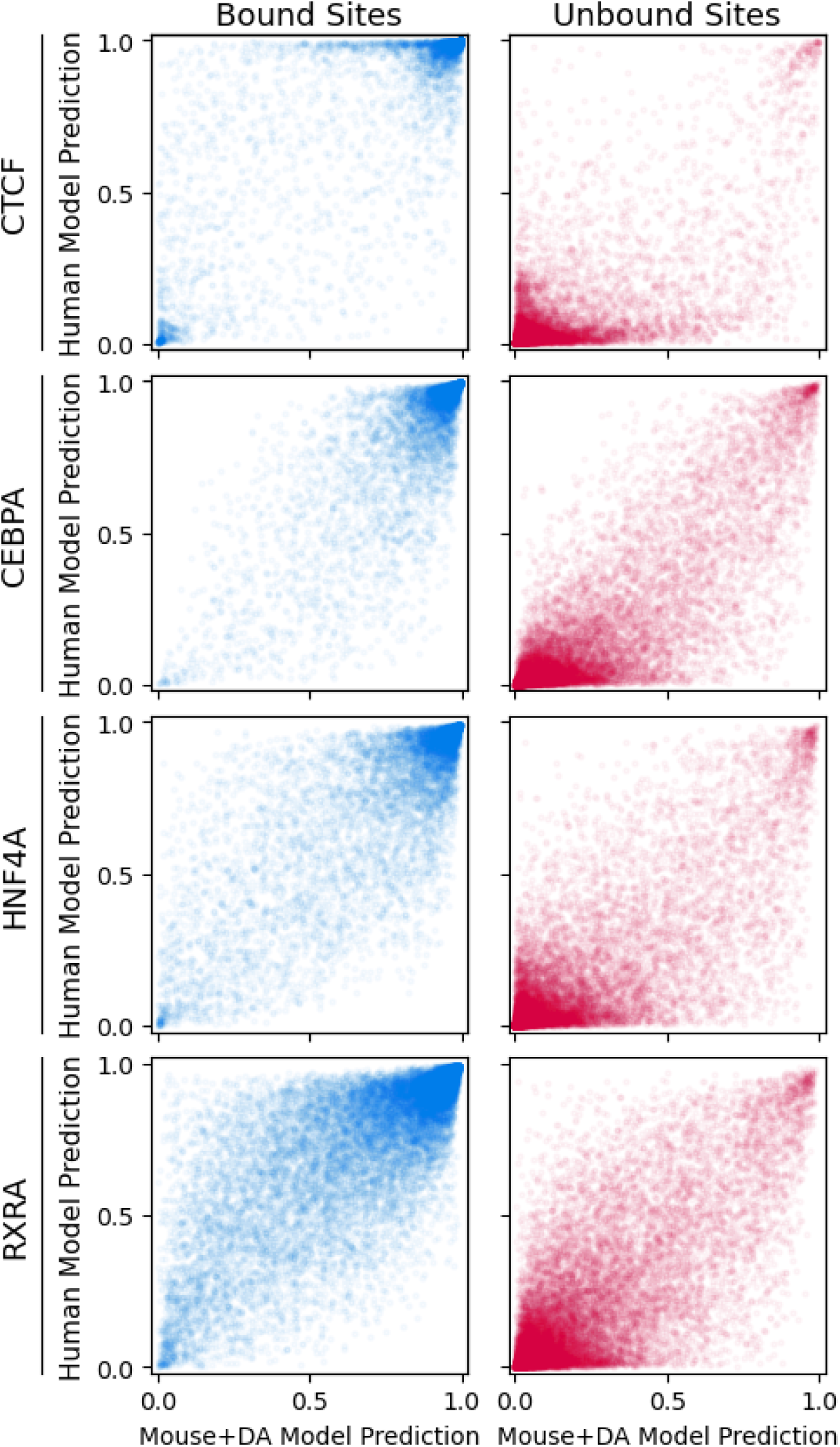
Differential predictions of human genome sites between human-trained and domain-adaptive mouse-trained models. Domain-adaptive mouse models, unlike the original mouse models, do not show species-specific systematic misprediction of unbound sites.

**Figure 11:**
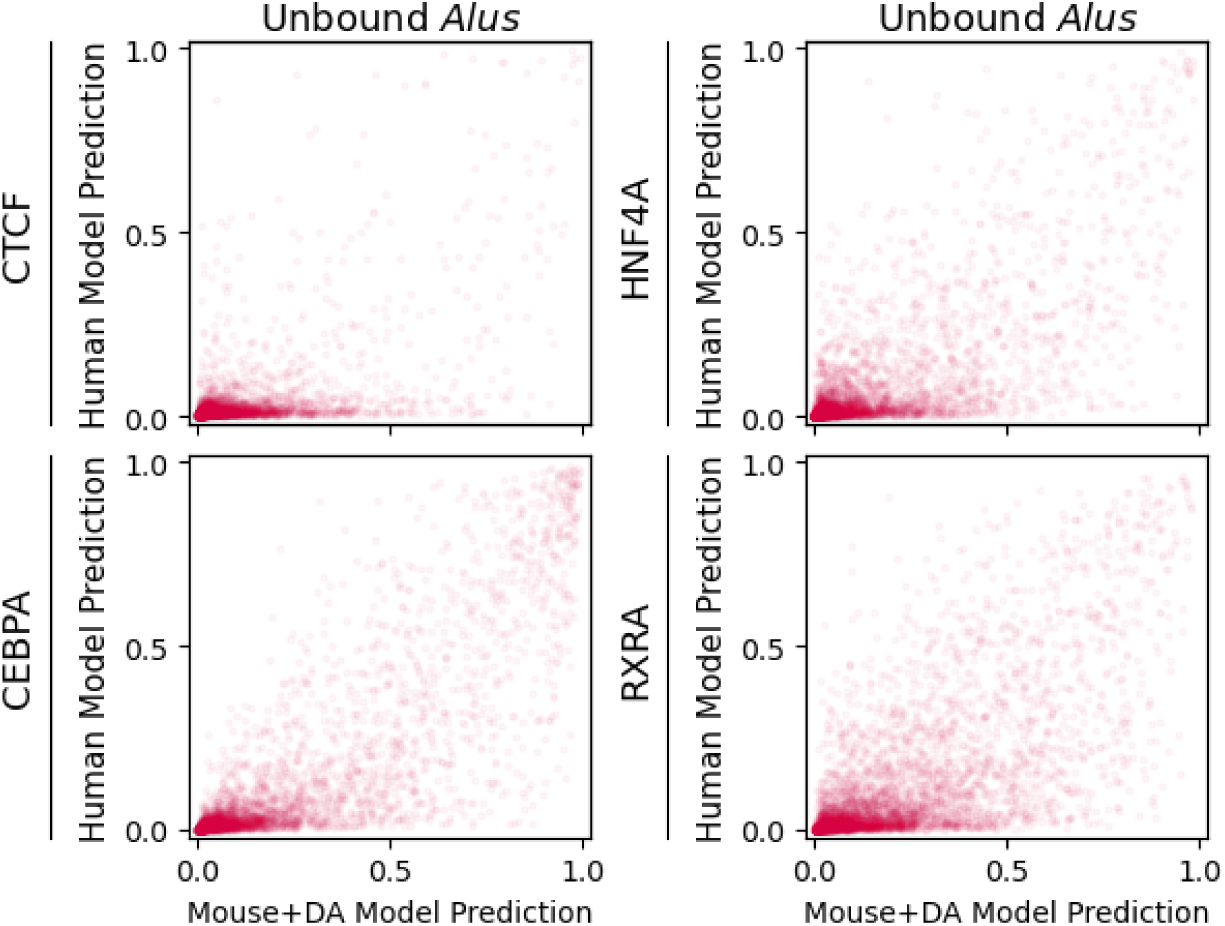
Differential predictions of unbound sites containing *Alu* elements between domain-adaptive mouse-trained models and human-trained models. Unlike the original mouse models, domain-adaptive mouse models do not show systematic overprediction of *Alu* repeats.

Compared to Figure 4, the mouse-model-specific false positives have diminished for all TFs. This suggests that the domain-adaptive models are able to correct the problem of false positive predictions from *Alus* by scoring unbound sites overlapping *Alus* lower than the basic model did. This effect is even present for CTCF, even though there was no noticeable auPRC difference for CTCF between domain-adaptive and basic mouse models – likely because the initial *Alu* enrichment in CTCF mouse-model false positives was lower than for other TFs.

In contrast, the site-distribution plots for bound sites demonstrate no noticeable difference from the original plots for the basic model architecture. We applied the same SeqUnwinder analysis to look for sequence features that discriminate between mouse-model false negatives and true positives and discovered similar, but not identical, motif-like short sequence patterns as we did previously (Figure S7). Thus, our domain adaptation approach does not appear to have any major influence on bound site predictions.

### *Alus* commonly drive mouse-model false positives across diverse cell types

Finally, we asked whether the observed over-prediction of species-specific repeats is a general issue of concern in cross-species TF binding prediction, or whether it is particular to the examined liver TFs. We thus widened our analyses to 53 additional pairs of ChIP-seq datasets targeting orthologous TFs across 8 additional equivalent human and mouse cell types (see Methods). One caveat is that the expanded set of paired datasets typically focus on cell lines and cell types that are more difficult to closely match across species than liver samples. Thus, the additional experiments examined here may not be as comparable across species as the previously examined liver datasets.

Our expanded analyses confirm that the crossspecies performance gap is present in most tested TFs and cell types (Table S3). A large portion of mouse-to-human false positive predictions is attributable to *Alu* elements. In 43 of the 53 additional examined datasets, *Alu* elements overlap a third or more of the mousemodel-unique false positive predictions (Table S4). Our domain adaptation procedure is successful in reducing *Alu*-related false positive predictions in 46 of the 53 additional examined datasets (Figure 12; Table S4). However, in megakaryocyte and hematopoietic progenitor datasets, we generally see a smaller percentage of mouse-model-unique false positives being attributable to *Alus*. The false positive predictions that do overlap *Alus* are also generally less likely to be corrected by our domain adaptation approach in these cell types (Figure 12). Therefore, our observations may not apply uniformly to all cell types.

**Figure 12:**
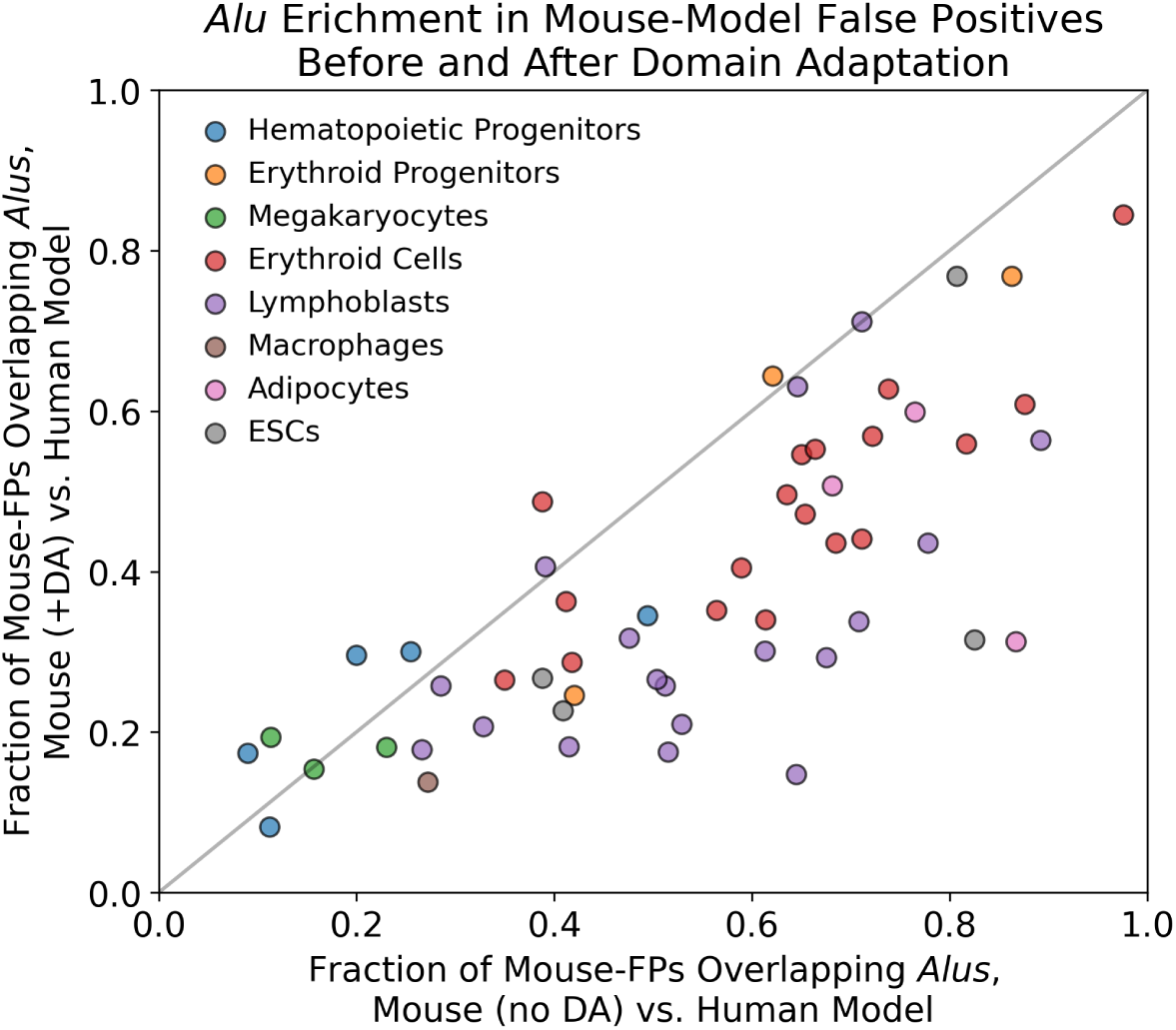
The fraction of mouse-model-unique false positives that overlap *Alus* when either the basic mouse model (x-axis) or the domain-adaptive mouse model (yaxis) are compared against the human model, across our additional paired datasets. The black diagonal line shows *y* = *x*; points below the line represent TFs where the fraction of *Alus* in mouse-model-unique false positives decreased with our domain adaptation strategy.

## Discussion

Enabling effective cross-species TF binding imputation strategies would be transformative for studying mammalian regulatory systems. For instance, TF binding information could be transferred from model organisms in cell types and developmental stages that are difficult or unethical to assay in humans. Similarly, one could annotate regulatory sites in non-model species of agricultural or evolutionary interest by leveraging the substantial investment that has been made to profile TF binding sites in human, mouse, and other model organisms (ENCODE Project Consortium 2012; Yue et al. 2014; Roadmap Epigenomics Consortium et al. 2015).

Our results suggest that cross-species TF binding imputation is feasible, but we also find a pervasive performance gap between within-species and cross-species prediction tasks. One set of culprits for this crossspecies performance gap are species-specific transposable elements. For example, models trained using mouse TF binding data have never seen an *Alu* SINE element during training, and often falsely predict that these elements are bound by the relevant TF. Since *Alu* elements appear at high frequency in the human genome, their misprediction constitutes a large proportion of the crossspecies false positive predictions, and thereby substantially affect the genome-wide performance metrics of the model. It should be noted that *Alus* and other transposable elements can serve as true regulatory elements (Bourque et al. 2008; Sundaram et al. 2014), and thus we don’t assume that all transposable elements should be labeled as TF “unbound”. Indeed, we minimized the potential mislabeling of truly bound transposable elements as “unbound” by focusing all our analyses on regions of the genome that have a high degree of mappability (and are thereby less likely to be subject to mappability-related false negative labeling issues in the TF ChIP-seq data).

We demonstrated that a simple domain adaptation approach is sufficient to correct the systematic mispredictions of *Alu* elements as TF bound. Training a parallel task (discriminating between species) but with gradient reversal employed during backpropagation has the effect of discouraging species-specific features being learned by the shared convolutional layers of the network. This approach is straightforward to implement and has the advantage that TF binding labels need only be known in the training species. Our approach accounts for domain shifts in the underlying genome sequence composition, assuming that the general features of TF binding sites are conserved within the same cell types across species.

We note that the underlying assumption of crossspecies TF binding prediction - i.e., that the overall features of cell-specific TF binding sites are conserved - may not hold true in all cases. Concordant importance scores between mouse and human models across true-positive bound sites suggests that both models learned similar representations of the TF’s cognate motif. However, we also observe that there are sequence features in bound sites that discriminate between correct and incorrect predictions specific to cross-species models. These discriminative sequence features suggest that cross-species false negative prediction errors could be the result of differential TF activity across the two species. Such differential activities could result from gain or loss of TF expression patterns, non-conserved cooperative binding capabilities, or evolved sequence preferences of the TF itself. We observe that these discriminative features are often preserved after we apply sequence composition domain adaptation, suggesting that our approach does not address the situation where TF binding logic is not fully conserved across species.

Other recent work has also demonstrated the feasibility of cross-species regulatory imputation. For example, Chen, et al. assessed the abilities of support vector machines (SVMs) and CNNs to predict potential enhancers (defined by combinations of histone marks) when trained and tested across species of varying evolutionary distances (Chen et al. 2018). Interestingly, they observed that while CNNs outperform SVMs in within-species enhancer prediction tasks, they are worse at generalizing across species. Our work suggests a possible reason for, and a solution to, this generalization gap. Two other recent manuscripts have applied more complex neural network architectures to impute TF binding and other regulatory signals across species (Kelley 2020; Schreiber et al. 2020). Those studies focus on models that are trained jointly across thousands of mouse and human regulatory genomic datasets. They thus assume that substantial amounts of regulatory information has already been characterized in the target species, which may not be true in some desired cross-species imputation settings. In general, however, joint modeling approaches are also likely to benefit from domain adaptation strategies that account for species-specific differences in sequence composition, and our results are thus complementary to these recent reports.

In summary, our work suggests that cross-species TF binding prediction approaches should beware of systematic differences between the compositions of training and test species genomes, including species-specific repetitive elements. Our contribution also suggests that domain adaptation is a promising strategy for addressing such differences and thereby making cross-species predictions more robust. Further work is needed to characterize additional sources of the cross-species performance gap and to generalize domain adaptation approaches to scenarios where training data is available from multiple species.

## Methods

### Data processing

Datasets were constructed by splitting the mouse (mm10) and human (hg38) genomes into 500 bp windows, offset by 50 bp. Any windows overlapping ENCODE blacklist regions were removed (Amemiya et al. 2019). We then calculated the fraction of each window that was uniquely mappable by 36 bp sequencing reads and retained only the windows that were at least 80% uniquely mappable (Karimzadeh et al. 2018). Mappability filtering was performed to remove potential peakcalling false negatives; otherwise, any genomic window too unmappable for confident peak-calling would be a potential false negative.

ChIP-seq experiments and corresponding controls (where available) were collected from ENCODE, GEO, and ArrayExpress. Database accession IDs for all data used in this study are listed in Tables S5, S6, and S7. We chose to focus our initial analyses on liver, as several previous studies have provided matched ChIP-seq experiments characterizing orthologous TF binding across mammalian liver samples (Schmidt et al. 2010; Odom et al. 2007). Our expanded analyses use erythroid, lymphoblast, and ES cell line experiments that were previously compared across species by Denas, et al. (Denas et al. 2015). We also analyzed matched adipocyte datasets that were performed on adipocyte cell lines within the same labs (Schmidt et al. 2011; Mikkelsen et al. 2010). Additional datasets were sourced by searching the literature for ChIP-seq data targeting orthologous TFs in erythroid progenitor, megakaryocyte, macrophage, and hematopoietic progenitor cell types (Tijssen et al. 2011; Hu et al. 2011; Pham et al. 2012; Pencovich et al. 2013; Kaikkonen et al. 2013; Beck et al. 2013; Yue et al. 2014; Huang et al. 2016; Goode et al. 2016).

For cell types where all data was sourced from the mouse and human ENCODE projects (i.e., erythroid, lymphoblast, and ES cell lines), we downloaded ChIP-seq narrow peak calls from the ENCODE portal. For liver and all other cell types, we first aligned the fastq files to the mm10 and hg38 reference genomes using bowtie (version 1.3.0) (Langmead and Salzberg 2012). We then called ChIP-seq peaks using MultiGPS v0.74 with default parameters, excluding ENCODE blacklist regions (Mahony et al. 2014; Amemiya et al. 2019). Corresponding control experiments were utilized during peak calling when available. Peak calls were converted to binary labels for each window in a genome: “bound” (1) if any peak center fell within the window, “unbound” (0) otherwise. Table S5 shows the numbers of peaks called for liver datasets, as well as the number of bound windows retained after filtering and the fraction of all retained windows that are bound; Tables S6 and S7 show the same information for all other datasets. Candidate datasets were discarded from the analysis if the numbers of called peaks was less than 1000 in mouse or human.

### Dataset splits for training and testing

Chromosomes 1 and 2 of both species were held out from all training datasets. For computational efficiency, one million randomly selected windows from Chromosome 1 were used as the validation set for each species (for hyperparameter tuning). All windows from Chromosome 2 were used as the test sets.

TF binding task training data was constructed identically for all model architectures. Since binary classifier neural networks often perform best when the classes are balanced in the training data, the binding task training dataset consisted of all bound examples and an equal number of randomly sampled (without replacement) unbound examples, excluding examples from Chromosomes 1 and 2. To increase the diversity of examples seen by the network across training, in each epoch a distinct random set of unbound examples was used, with no repeated unbound examples across epochs.

Domain-adaptive models also require an additional “species-background” training set from both species for the species discrimination task. Species-background data consisted of randomly selected (without replacement) examples from all chromosomes except 1 and 2. Binding labels were not used in the construction of these training sets. In each batch, the species-background examples were balanced, with 50% human and 50% mouse examples, and labeled according to their species of origin (not by binding). The total number of species-background examples in each batch was double the number of binding examples.

### Basic model architecture

The network takes in a one-hot encoded 500 bp window of DNA sequence and passes it through a convolutional layer with 240 20-bp filters, followed by a ReLU activation and max-pooling (pool window and stride of 15 bp). After the convolutional layer is an LSTM with 32 internal nodes, followed by a 1024-neuron fully-connected layer with ReLU activation, followed by a 50% Dropout layer, followed by a 512-neuron fully-connected layer with sigmoid activation. The final layer is a single sigmoid-activated neuron.

### Domain-adaptive model architecture

The domain-adaptive network builds upon the basic model described above by adding a new “species discriminator” task. The network splits into two output halves following max-pooling after the convolutional layer. The max-pooling output feeds into a gradient reversal layer (GRL) – the GRL merely outputs the identity of its input during the feed-forward step of model training, but during backpropagation, it multiplies the gradient of the loss by −1. The GRL is followed by a Flatten layer, a ReLU-activated fully connected layer with 1024 neurons, a sigmoid-activated fully connected layer of 512 neurons, and finally a single-neuron layer with sigmoid activation.

### Model training

All models were trained with Keras v2.3.1 using the Adam optimizer with default parameters (Chollet 2015; Kingma and Ba 2014). Training ran for 15 epochs, with models saved after each epoch. After training, we selected models for downstream analysis by choosing the saved model with highest auPRC on the training-species validation set.

The basic models were trained by standard procedure with a batch size of 400 (see Section 2.1.2 for training dataset construction). The domain-adaptive models, on the other hand, required a more complex batching setup. Because domain-adaptive models predict two tasks – binding and the species of origin of the input sequence – they require two stages of dataset input per batch. The first stage is identical to a basic model training batch, but with ⌊400*/*3⌋ = 133 binding examples from the source species. The second stage uses ⌈400 ∗ 2*/*3 ⌉ = 267 examples each from the source species’ and target species’ “species-background” datasets.

Crucially, the stages differ in how task labels are masked. For each stage, only one of the two output halves of the network trains (the loss backpropagates from one output only). In the first stage, we mask the species discriminator task, so that only the binding task half of the model trains on binding examples from the training species. In the second stage, we mask the binding task, so only the species discriminator task half trains. Thus, the binding task only trains on examples from the source species, while the species discriminator task doesn’t see binding labels from either species.

Meanwhile, the weights of the shared convolutional layer are influenced by both tasks. Because these stages occur within a single batch and not in alternating batches, they concurrently influence the weights of the convolutional filters; there is no oscillating “back-and-forth” between the two tasks from batch to batch.

### Differentially-predicted site categorization

To quantify site enrichment within discrete categories such as “false positives” and “false negatives”, it was necessary to define the boundaries for these labels. In particular, when comparing prediction distributions between models, we needed to define what constitutes, for instance, a “false positive unique to model *A*.” We constructed the following rules for site categorization: 1) unbound sites must have predictions above 0.5 to be labeled false positives, and bound sites must have predictions below 0.5 to be labeled false negatives; 2) a site is considered to be differentially predicted between two source species *A* and *B* if |*P*_*A*_ − *P*_*B*_| *>* 0.5, where *P*_*A*_ and *P*_*B*_ are the predictions from models trained on data from species *A* and species *B*, respectively; 3) only sites meeting this differential prediction threshold are labeled as a false positive or negative unique to one model. Thus, if we are comparing models from species *A* and *B*, and a site is labeled a false positive unique to model *A*, then *P*_*A*_ *>* 0.5 and *P*_*B*_ < 0.5. To reduce noise in these categorizations, rather than letting *P*_*A*_ and *P*_*B*_ equal the predictions from single models, we trained 5 independent replicate models for each TF and source species, and then let *P*_*A*_ be the average prediction across the 5 replicate models trained on data from species *A* for a given TF.

### Bound site discriminative motif discovery

SeqUnwinder (v. 0.1.3) (Kakumanu et al. 2017) was used to find motifs that discriminate between true positive predictions and mouse-model-specific false negative predictions using the following command-line settings: “--threads 10 --makerandregs --makerandregs - -win 500 --mink 4 --maxk 5 --r 10 --x 3 --a 400 -hillsthresh 0.1 --memesearchwin 16”, and using MEME v. 5.1.0 (Machanick and Bailey 2011) internally.

### Repeat analysis

All repeat analysis used the RepeatMasker track from the UCSC Genome Browser (Smit et al. 1996). Genome windows were labeled as containing an *Alu* element if there was any overlap (1 or more bp) with any *Alu* annotation. For Table S1, repeat classes were excluded if fewer than 500 examples of that class were annotated in the test chromosome (before mappability filtering).

### Gapped *k*-mer SVMs

The gkmtrain and gkmpredict utilities from the lsgkm package were used for gkmSVMs gkm training and prediction generation, respectively (Lee 2016). For training, 50000 examples each were selected randomly from the set of all bound windows and unbound windows in the original deep learning model training sets. Every 10th example from the original test set (in other words, sampling windows such that all selected windows were nonoverlapping) was considered in evaluation for computational efficiency. All default parameters were used in running lsgkm (center-weighted + truncated *l*-mer kernel, word length 11, maximum 3 mismatches).

### Importance scoring

For a given 500bp window and model, importance scores were generated using a method similar to in silico mutagenesis, which measures the change in model prediction when a given base and the region immediately around it are ablated. First, ten independent dinucleotide-shuffled versions of the original sequence were generated to serve as reference sequences unlikely to contain motifs. Next, the 5bp region centered at a particular base was replaced with the corresponding 5bp region from one of the ten shuffled sequences, and the post-sigmoid difference in model output for this ablated sequence was recorded. This was repeated for all ten shuffled sequences, with the average model prediction differential reported as the score for the base that the ablated region centered on. This process was repeated for all bases in the sequence being scored.

## Software availability

Open source code (MIT license) is available from: https://github.com/seqcode/cross-species-domain-adaptation

## Acknowledgements

The authors thank the members of the Center for Eukaryotic Gene Regulation at Penn State and Jacob Schreiber for helpful feedback and discussion.

---

**Figure S1:**
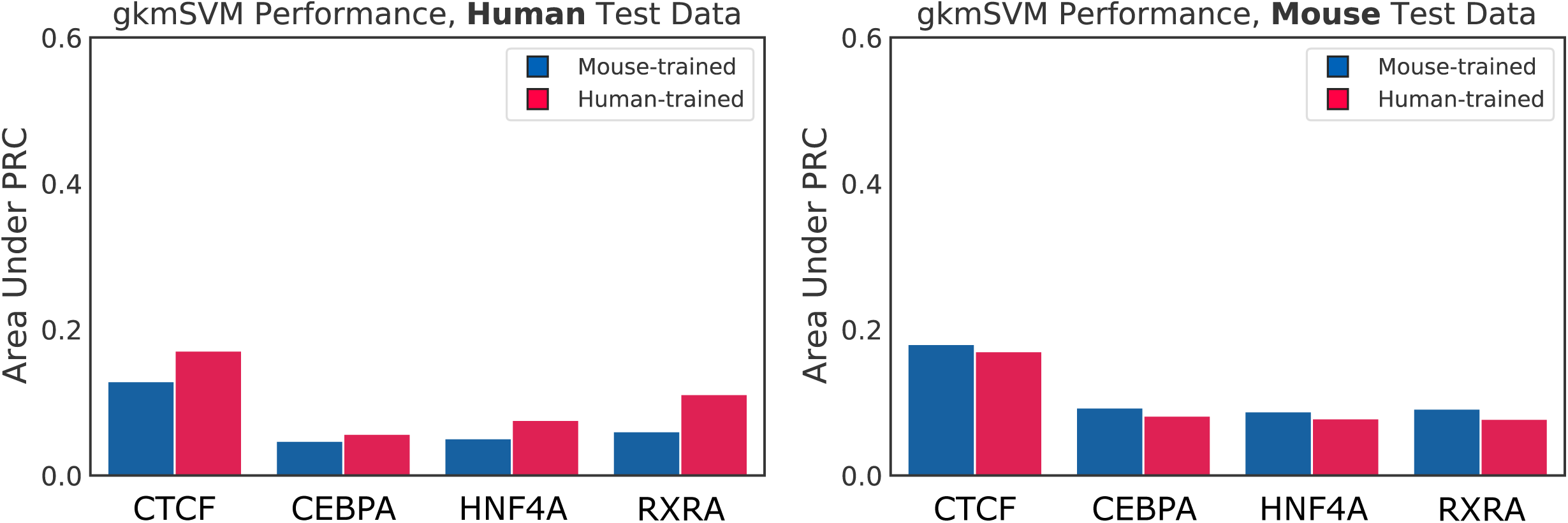
Results of evaluating the performance of mouse-trained (blue) and human-trained (red) gapped *k*-mer SVM models on non-overlapping windows from the human (left) and mouse (right) test datasets (Chromosome 2). For each TF and species, an SVM was trained using a balanced set of bound and unbound windows from the original training set.

**Figure S2:**
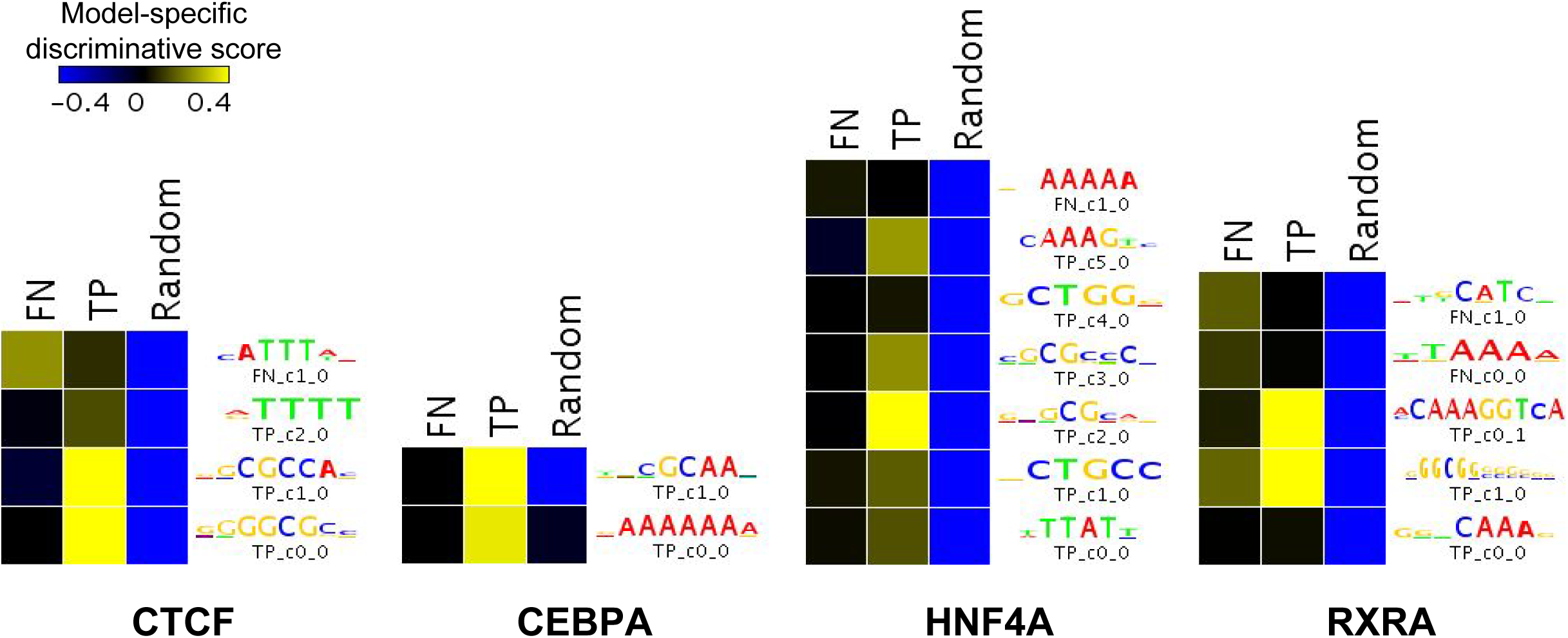
Motif-like sequence features can discriminate between human-genome bound sites correctly predicted by mouse-trained and human-trained models (true positives or TP) and bound sites correctly predicted only by human-trained models (mouse-specific false negatives or FN) for each TF. See Methods for site categorization details.

**Figure S3:**
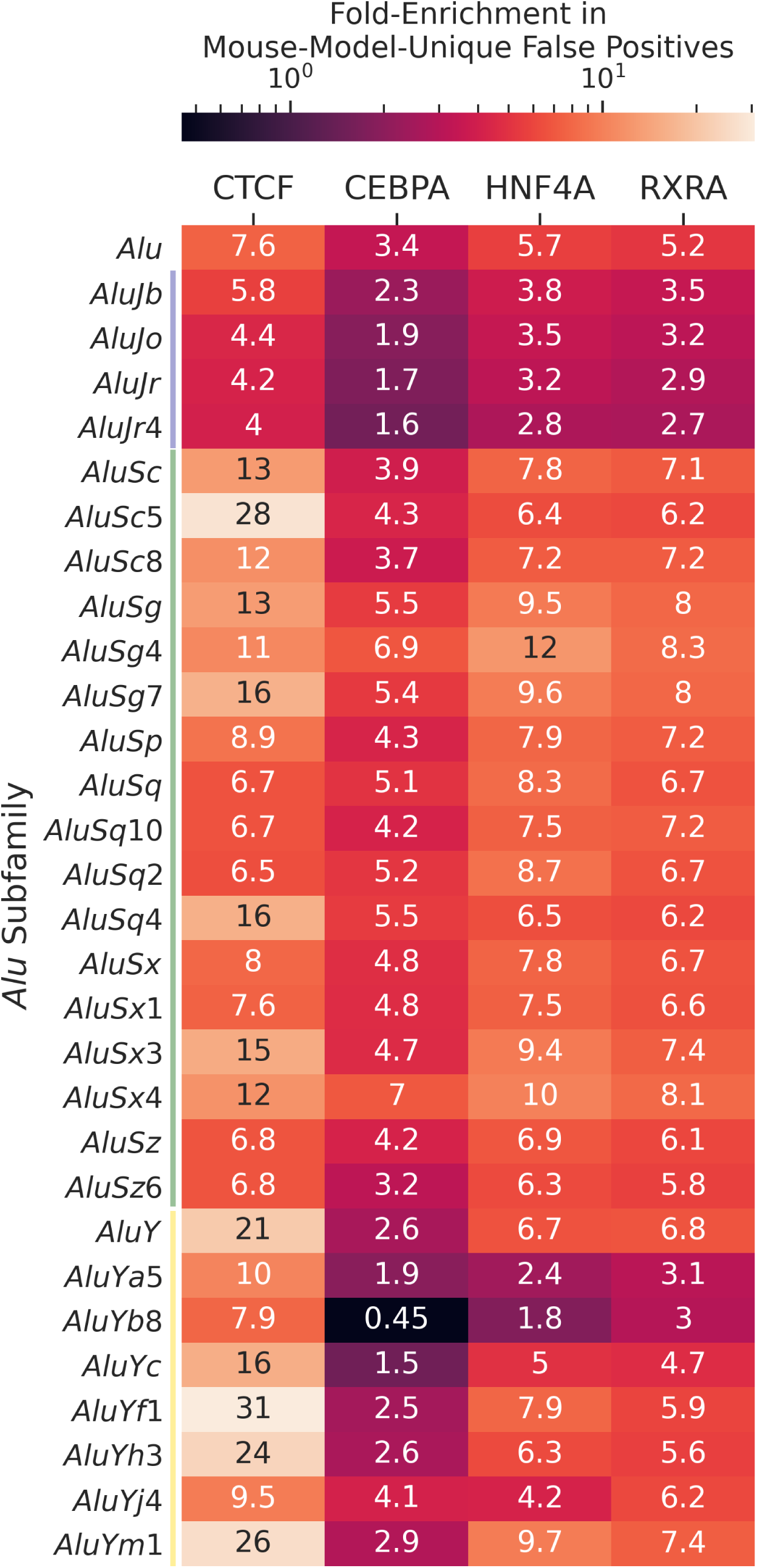
Enrichment of specific *Alu* subfamilies within the set of false positives unique to the mouse-trained model, relative to false positives common to both mouse-trained and human-trained models. For a given TF and *Alu* subfamily, the fraction of windows overlapping any RepeatMasker-annotated instance of that repeat type were calculated for both classes of false positives. The values in the figure show the ratio between *Alu* fractions in mousemodel-unique false positives and both-model false positives. Only *Alu* subamilies with at least 100 annotated examples in the test dataset (Chromosome 2) and covering a non-zero fraction (at least 0.01%) of both false positive categories are included.

**Figure S4:**
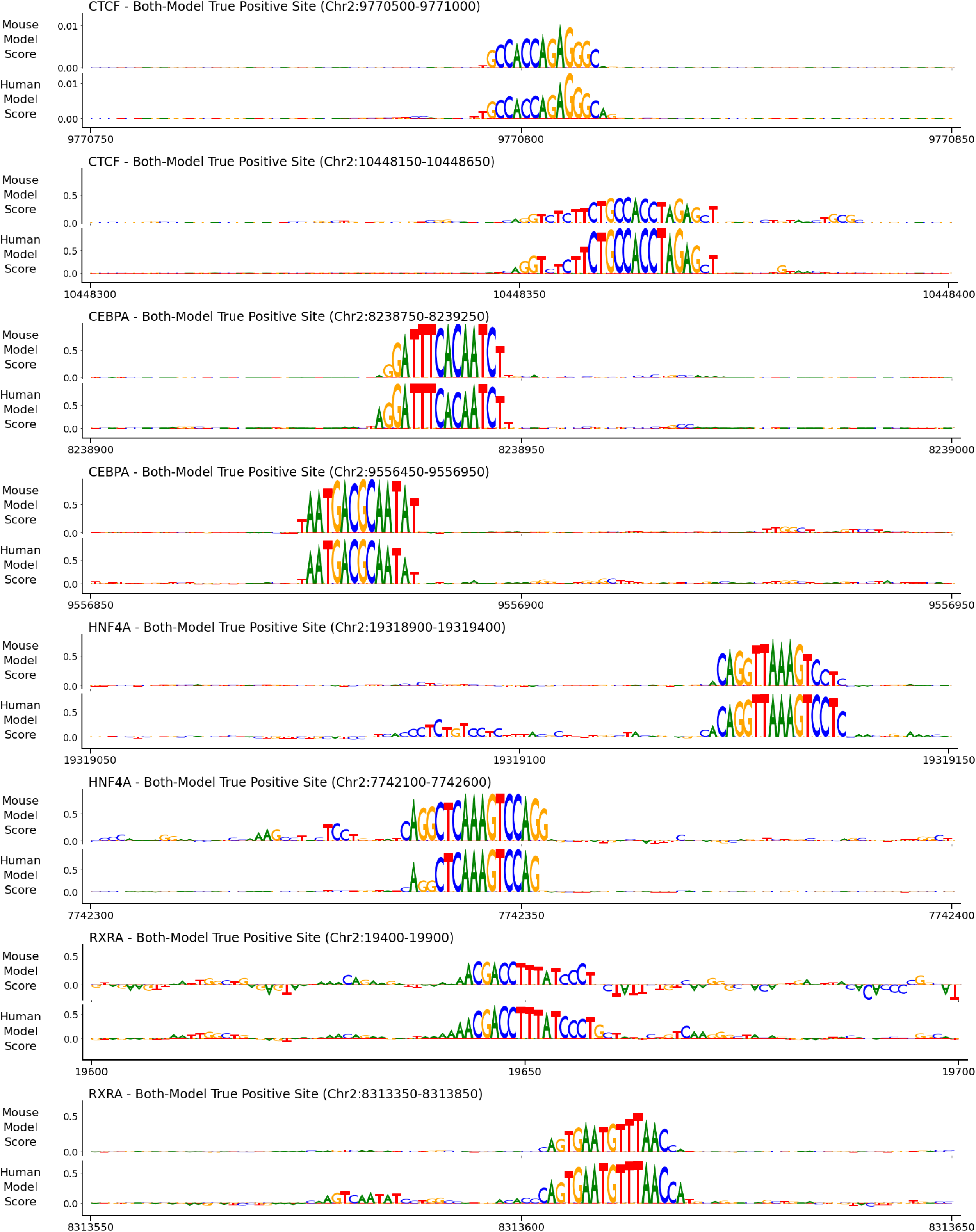
Importance scores for example both-model true positive sites for the 4 TFs. Bases were scored using a modified ISM algorithm (see Methods). The 500bp example sites have been enlarged and cropped around motif-like instances for readability. 19

**Figure S5:**
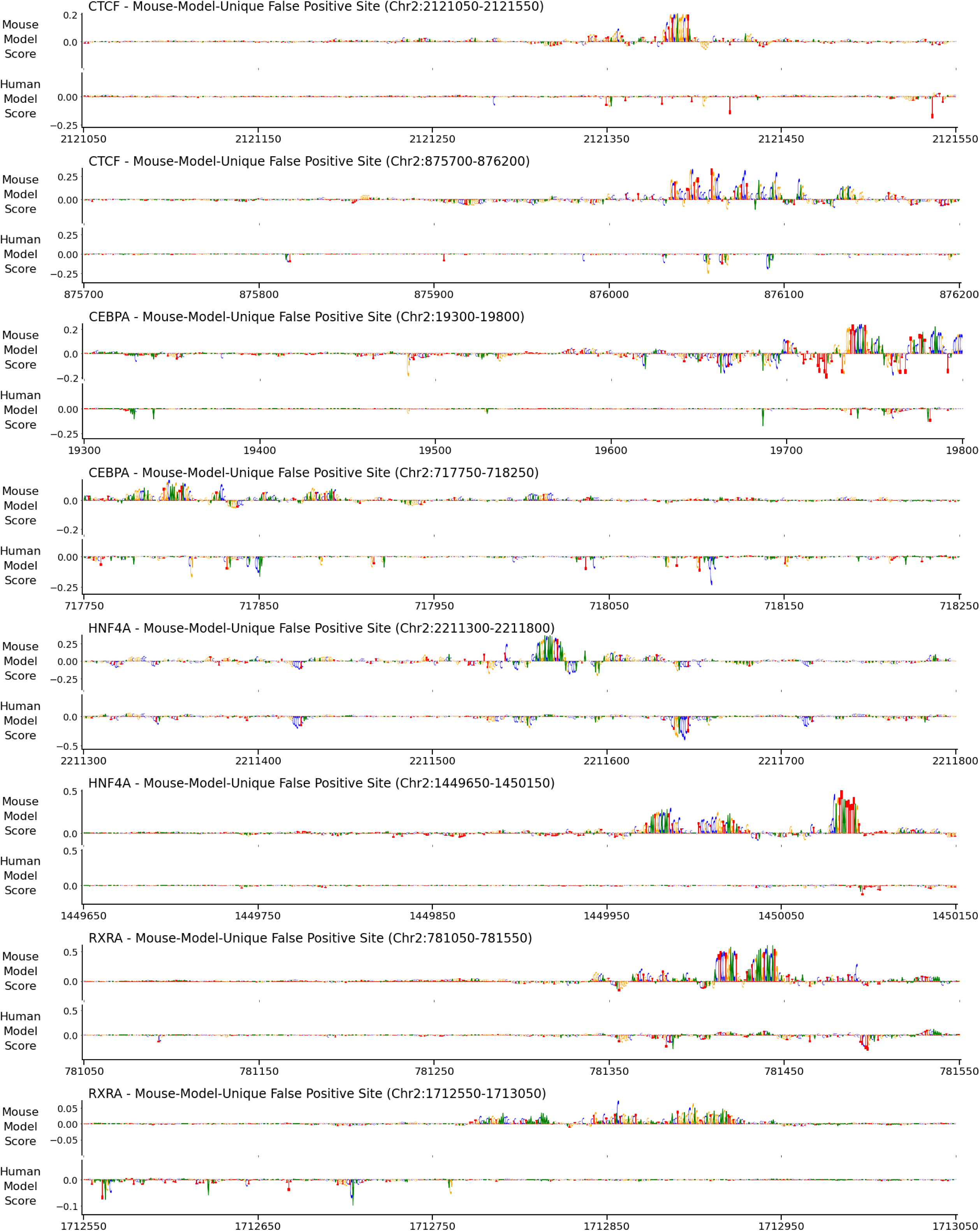
Importance scores for example false positive sites mispredicted only by the mouse model. Bases were scored using a modified ISM algorithm (see Methods). 20

**Figure S6:**
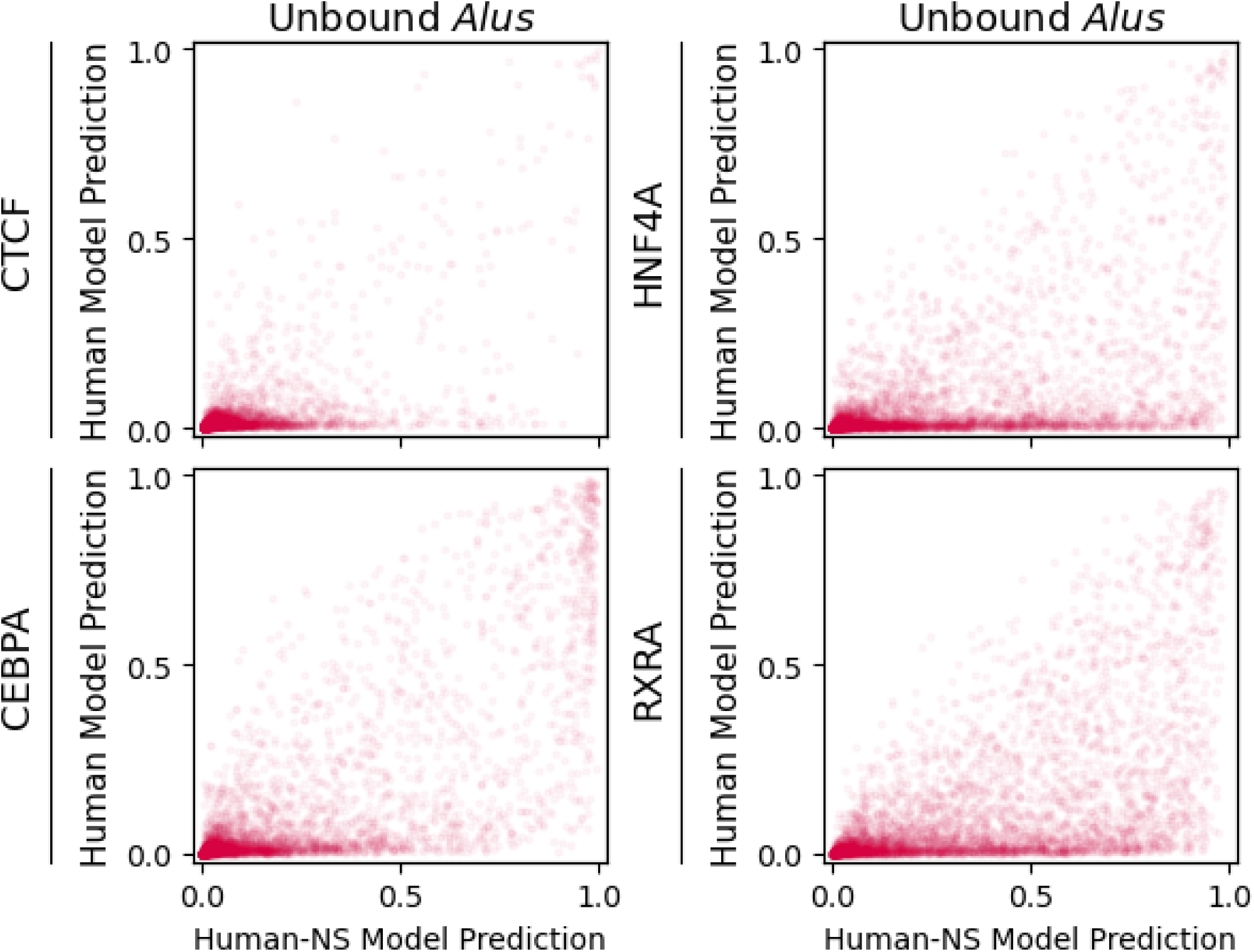
Comparison between the predictions of human-trained models that were trained without examples overlapping SINEs (x-axis) to the predictions of standard human-trained models (y-axis). Unbound *Alu* repeats make up a large part of the false positives unique to the no-SINEs model. For visual clarity, only 5% of windows are shown.

**Figure S7:**
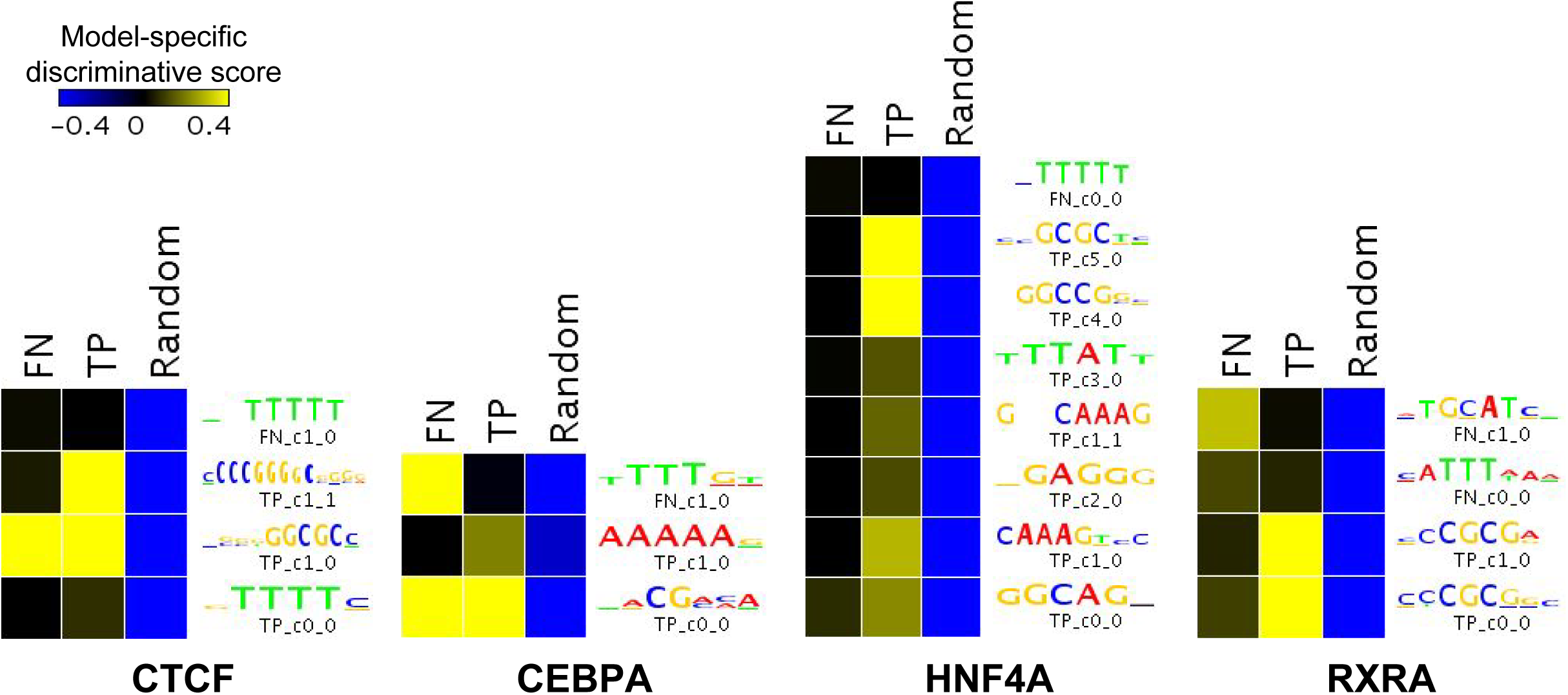
False negative predictions unique to mouse-trained models trained with domain adaptation, compared to human-trained models, can be distinguished from true positive predictions through motif-like sequence features. See Methods for site categorization details.

**Table S1:** Percent of windows overlapping various RepeatMasker-defined repeat elements, for different categories of genomic windows from the held-out test set. Only RepeatMasker repeat classes with at least 500 distinct annotations within the test set are shown. FPs: false positives. FNs: false negatives. Mouse Only: specific to mouse-trained models. See Methods for more details on site categorization.

| TF | Bound | FN (Both Models) | FN (Mouse Only) | Unbound | FP (Both Models) | FP (Mouse Only) |
| --- | --- | --- | --- | --- | --- | --- |
| **DNA** |  |  |  |  |  |  |
| CTCF | 10.1% | 11.6% | 12.0% | 11.4% | 7.2% | 7.9% |
| CEBPA | 12.3% | 11.0% | 0.0% | 11.4% | 13.0% | 9.5% |
| HNF4A | 10.8% | 11.4% | 11.6% | 11.4% | 9.8% | 9.0% |
| RXRA | 10.1% | 11.1% | 11.0% | 11.4% | 10.2% | 9.4% |
| **LINE** |  |  |  |  |  |  |
| CTCF | 18.0% | 22.2% | 24.9% | 37.6% | 14.6% | 24.8% |
| CEBPA | 25.5% | 25.7% | 0.0% | 37.6% | 29.0% | 32.3% |
| HNF4A | 21.0% | 25.4% | 30.7% | 37.6% | 21.8% | 29.7% |
| RXRA | 20.8% | 28.3% | 21.4% | 37.8% | 21.8% | 33.1% |
| **Low-Complexity** |  |  |  |  |  |  |
| CTCF | 2.8% | 2.2% | 3.0% | 1.9% | 4.5% | 1.5% |
| CEBPA | 1.5% | 1.6% | 0.0% | 1.9% | 1.6% | 1.4% |
| HNF4A | 2.0% | 1.4% | 1.5% | 1.9% | 2.4% | 1.5% |
| RXRA | 2.0% | 1.4% | 1.7% | 1.9% | 2.2% | 1.5% |
| **LTR** |  |  |  |  |  |  |
| CTCF | 8.7% | 12.6% | 7.1% | 17.6% | 13.5% | 13.6% |
| CEBPA | 12.8% | 11.1% | 0.0% | 17.6% | 19.0% | 14.6% |
| HNF4A | 13.3% | 14.9% | 11.4% | 17.6% | 19.5% | 13.9% |
| RXRA | 11.8% | 13.9% | 10.9% | 17.6% | 18.0% | 11.7% |
| **Simple Repeat** |  |  |  |  |  |  |
| CTCF | 13.9% | 11.2% | 8.1% | 11.5% | 17.1% | 10.6% |
| CEBPA | 9.4% | 9.6% | 40.0% | 11.6% | 9.9% | 11.7% |
| HNF4A | 12.4% | 12.0% | 11.2% | 11.5% | 11.5% | 12.1% |
| RXRA | 11.5% | 9.7% | 14.4% | 11.5% | 11.1% | 13.4% |
| **SINE** |  |  |  |  |  |  |
| CTCF | 23.5% | 25.1% | 20.4% | 31.2% | 16.8% | 68.9% |
| CEBPA | 30.8% | 22.5% | 0.0% | 31.1% | 35.5% | 84.8% |
| HNF4A | 27.1% | 21.8% | 20.5% | 31.2% | 28.6% | 88.7% |
| RXRA | 26.9% | 24.2% | 17.5% | 31.2% | 31.4% | 97.1% |
| **Unknown** |  |  |  |  |  |  |
| CTCF | 0.2% | 0.0% | 0.2% | 0.2% | 0.1% | 0.0% |
| CEBPA | 0.3% | 0.5% | 0.0% | 0.2% | 0.2% | 0.1% |
| HNF4A | 0.2% | 0.1% | 0.2% | 0.2% | 0.2% | 0.0% |
| RXRA | 0.2% | 0.2% | 0.1% | 0.1% | 0.2% | 0.0% |

**Table S2:** Percent of windows overlapping an *Alu* element when domain-adaptive mouse models are compared to human models (compare to Table 1). The fraction of mouse-model-unique false positives overlapping *Alu* elements (right-most column) have decreased drastically for all TFs. FPs: false positives. FNs: false negatives.

| TF | Bound | FNs (Both Models) | FNs (Mouse Only) | Unbound | FPs (Both Models) | FPs (Mouse Only) |
| --- | --- | --- | --- | --- | --- | --- |
| CTCF | 12.1% | 11.8% | 12.7% | 21.2% | 8.2% | **21.0%** |
| CEBPA | 18.3% | 15.8% | 20.0% | 21.2% | 23.2% | **43.2%** |
| HNF4A | 13.6% | 14.9% | 14.5% | 21.2% | 14.2% | **29.4%** |
| RXRA | 13.3% | 15.8% | 10.0% | 21.3% | 15.2% | **51.7%** |

**Table S3:** Average auPRC values from evaluating the basic mouse models, domain-adaptive mouse models, and basic human models on the mouse (left columns) and human (right columns) test sets, across all additional datasets beyond the primary liver TFs. The auPRCs shown are the average across three replicate model trainings for basic mouse-trained and human-trained models and across two replicate model trainings for domain-adaptive mouse models. Note that because the auPRC metric depends on the sparsity of the positive class (bound sites), these values are not comparable across test sets, across TFs, or across cell types.

| TF | auPRC, Mouse Test Set |  |  | auPRC, Human Test Set |  |  |
| --- | --- | --- | --- | --- | --- | --- |
|  | Mouse(Basic) | Mouse(+DA) | Human | Mouse(Basic) | Mouse(+DA) | Human |
| **Adipocytes** |  |  |  |  |  |  |
| CEBPA | 0.18 | 0.18 | 0.17 | 0.18 | 0.21 | 0.35 |
| CTCF | 0.67 | 0.66 | 0.55 | 0.56 | 0.56 | 0.62 |
| PPARG | 0.08 | 0.07 | 0.08 | 0.07 | 0.06 | 0.22 |
| **Erythroid Cells** |  |  |  |  |  |  |
| BHLHE40 | 0.09 | 0.09 | 0.07 | 0.13 | 0.13 | 0.19 |
| CTCF | 0.71 | 0.68 | 0.62 | 0.60 | 0.58 | 0.67 |
| E2F4 | 0.10 | 0.07 | 0.09 | 0.17 | 0.17 | 0.23 |
| ELF1 | 0.28 | 0.28 | 0.27 | 0.26 | 0.26 | 0.34 |
| ETS1 | 0.16 | 0.16 | 0.05 | 0.11 | 0.10 | 0.21 |
| GATA1 | 0.20 | 0.19 | 0.11 | 0.09 | 0.09 | 0.10 |
| JUND | 0.05 | 0.03 | 0.02 | 0.10 | 0.09 | 0.26 |
| MAFK | 0.14 | 0.12 | 0.14 | 0.17 | 0.16 | 0.39 |
| MAX | 0.18 | 0.18 | 0.14 | 0.19 | 0.20 | 0.26 |
| MAZ | 0.15 | 0.15 | 0.14 | 0.21 | 0.22 | 0.32 |
| MEF2A | 0.03 | 0.01 | 0.02 | 0.02 | 0.01 | 0.04 |
| MXI1 | 0.20 | 0.21 | 0.16 | 0.09 | 0.10 | 0.10 |
| MYC | 0.14 | 0.14 | 0.09 | 0.17 | 0.18 | 0.23 |
| NRF1 | 0.33 | 0.32 | 0.22 | 0.33 | 0.35 | 0.36 |
| TAL1 | 0.14 | 0.14 | 0.11 | 0.14 | 0.14 | 0.19 |
| UBTF | 0.15 | 0.15 | 0.15 | 0.19 | 0.19 | 0.23 |
| USF1 | 0.21 | 0.18 | 0.16 | 0.17 | 0.16 | 0.25 |
| USF2 | 0.12 | 0.11 | 0.09 | 0.14 | 0.16 | 0.13 |
| **Erythroid Progenitors** |  |  |  |  |  |  |
| CTCF | 0.69 | 0.67 | 0.57 | 0.60 | 0.59 | 0.67 |
| GATA1 | 0.09 | 0.09 | 0.08 | 0.10 | 0.08 | 0.16 |
| TAL1 | 0.06 | 0.04 | 0.07 | 0.08 | 0.07 | 0.21 |
| **ESCs** |  |  |  |  |  |  |
| CTCF | 0.78 | 0.76 | 0.71 | 0.53 | 0.54 | 0.66 |
| MAFK | 0.43 | 0.40 | 0.40 | 0.31 | 0.28 | 0.34 |
| NANOG | 0.14 | 0.12 | 0.05 | 0.05 | 0.05 | 0.08 |
| POU5F1 | 0.11 | 0.10 | 0.09 | 0.07 | 0.06 | 0.09 |
| **Hematopoietic Progenitors** |  |  |  |  |  |  |
| FLI1 | 0.21 | 0.16 | 0.09 | 0.06 | 0.06 | 0.17 |
| LMO2 | 0.06 | 0.04 | 0.00 | 0.00 | 0.00 | 0.01 |
| RUNX1 | 0.06 | 0.04 | 0.05 | 0.05 | 0.05 | 0.20 |
| SPI1 | 0.32 | 0.28 | 0.32 | 0.38 | 0.38 | 0.62 |

Table S3: Average auPRC values from evaluating the basic mouse models, domain-adaptive mouse models, and basic human models on the mouse (left columns) and human (right columns) test sets, across all additional datasets beyond the primary liver TFs.
| TF | auPRC, Mouse Test Set |  |  | auPRC, Human Test Set |  |  |
| --- | --- | --- | --- | --- | --- | --- |
|  | Mouse(Basic) | Mouse(+DA) | Human | Mouse(Basic) | Mouse(+DA) | Human |
| **Lymphoblasts** |  |  |  |  |  |  |
| BHLHE40 | 0.23 | 0.21 | 0.15 | 0.13 | 0.14 | 0.17 |
| CTCF | 0.70 | 0.69 | 0.58 | 0.63 | 0.61 | 0.65 |
| E2F4 | 0.12 | 0.09 | 0.12 | 0.12 | 0.11 | 0.13 |
| ELF1 | 0.32 | 0.30 | 0.27 | 0.34 | 0.34 | 0.34 |
| ETS1 | 0.16 | 0.15 | 0.05 | 0.05 | 0.05 | 0.19 |
| IRF4 | 0.23 | 0.22 | 0.14 | 0.11 | 0.10 | 0.14 |
| JUND | 0.09 | 0.07 | 0.05 | 0.04 | 0.04 | 0.07 |
| MAX | 0.17 | 0.17 | 0.13 | 0.17 | 0.18 | 0.19 |
| MAZ | 0.13 | 0.12 | 0.12 | 0.20 | 0.20 | 0.24 |
| MEF2A | 0.16 | 0.14 | 0.09 | 0.06 | 0.06 | 0.11 |
| MXI1 | 0.19 | 0.20 | 0.18 | 0.14 | 0.15 | 0.16 |
| MYC | 0.14 | 0.14 | 0.07 | 0.08 | 0.10 | 0.11 |
| NRF1 | 0.32 | 0.30 | 0.25 | 0.38 | 0.34 | 0.45 |
| TBP | 0.16 | 0.15 | 0.14 | 0.09 | 0.09 | 0.11 |
| TCF12 | 0.24 | 0.23 | 0.17 | 0.12 | 0.11 | 0.14 |
| USF1 | 0.22 | 0.20 | 0.17 | 0.20 | 0.20 | 0.19 |
| USF2 | 0.16 | 0.15 | 0.12 | 0.10 | 0.10 | 0.09 |
| **Macrophages** |  |  |  |  |  |  |
| SPI1 | 0.41 | 0.41 | 0.33 | 0.29 | 0.30 | 0.46 |
| **Megakaryocytes** |  |  |  |  |  |  |
| FLI1 | 0.26 | 0.15 | 0.22 | 0.15 | 0.07 | 0.16 |
| GATA1 | 0.09 | 0.08 | 0.02 | 0.03 | 0.02 | 0.04 |
| RUNX1 | 0.08 | 0.06 | 0.04 | 0.13 | 0.12 | 0.28 |

**Table S4:**
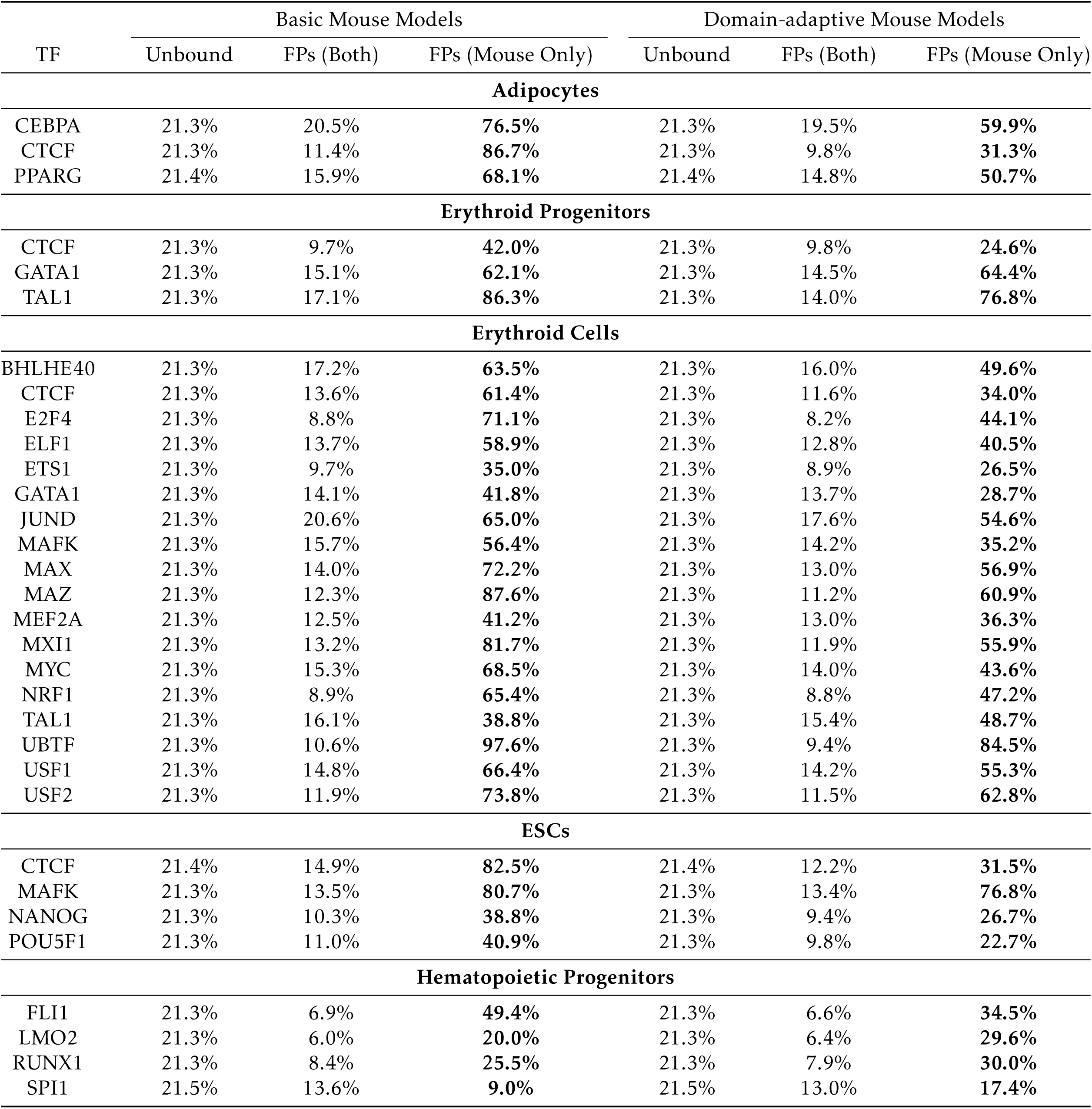
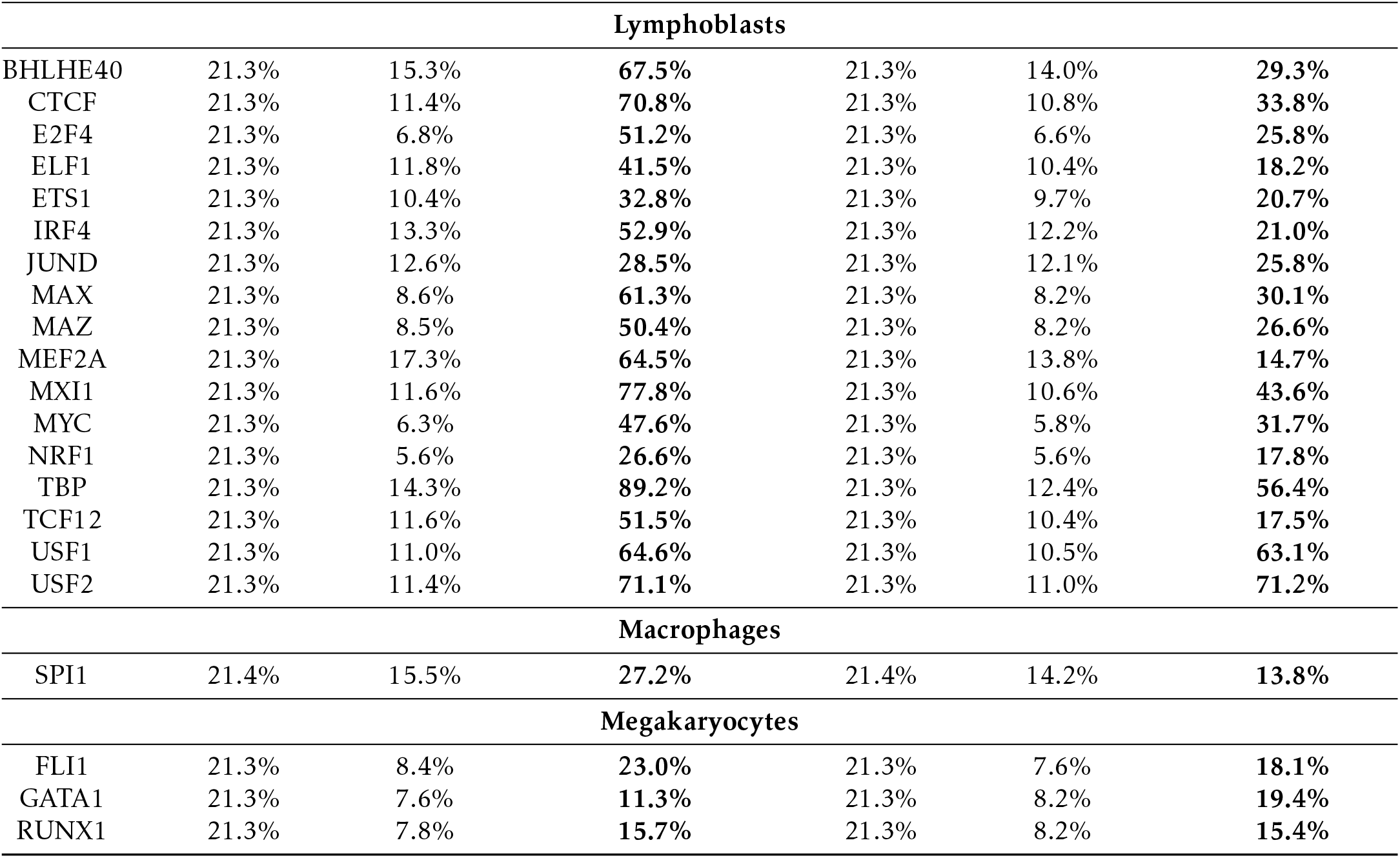
The percent of sites overlapping an *Alu* element without (left column set) or with domain adaptation (right column set), for each of the additional datsets included in Figure 12. FPs: false positives; either the set of unbound sites mispredicted as bound by both the mouse model and the human model, or false positives mispredicted by the mouse model only. See Methods for site categorization details.

**Table S5:** For the primary experimental data used in this study, the following quantities are listed: the number of peaks called across the entire genome; the number of called peaks within the filtered window set, merged if within 500 bp of each other; the number of windows in the filtered window set labeled bound due to peak overlap; the fraction of the filtered window set labeled bound; and the database accession ID (ENCODE, GEO, or ArrayExpress). The size of the filtered window sets for the mouse and human genomes were 44288170 and 51548966, respectively.

| TF | Species | Raw Peaks | Filtered Peaks | Bound Windows | Frac. Bound | Accession ID |
| --- | --- | --- | --- | --- | --- | --- |
| CTCF | Mouse | 34362 | 29677 | 317396 | 0.72% | ENCSR000CBU |
|  | Human | 34921 | 31812 | 330941 | 0.64% | GSE105829 |
| CEBPA | Mouse | 62636 | 49338 | 576787 | 1.30% | E-TABM-722 |
|  | Human | 32243 | 29244 | 306321 | 0.59% | E-TABM-722 |
| HNF4A | Mouse | 44800 | 36805 | 421268 | 0.95% | E-TABM-722 |
|  | Human | 42766 | 35352 | 396294 | 0.77% | E-TABM-722 |
| RXRA | Mouse | 50959 | 35221 | 446830 | 1.01% | GSM1299600 |
|  | Human | 93895 | 69999 | 863861 | 1.68% | ENCSR098XMN |

**Table S6:**
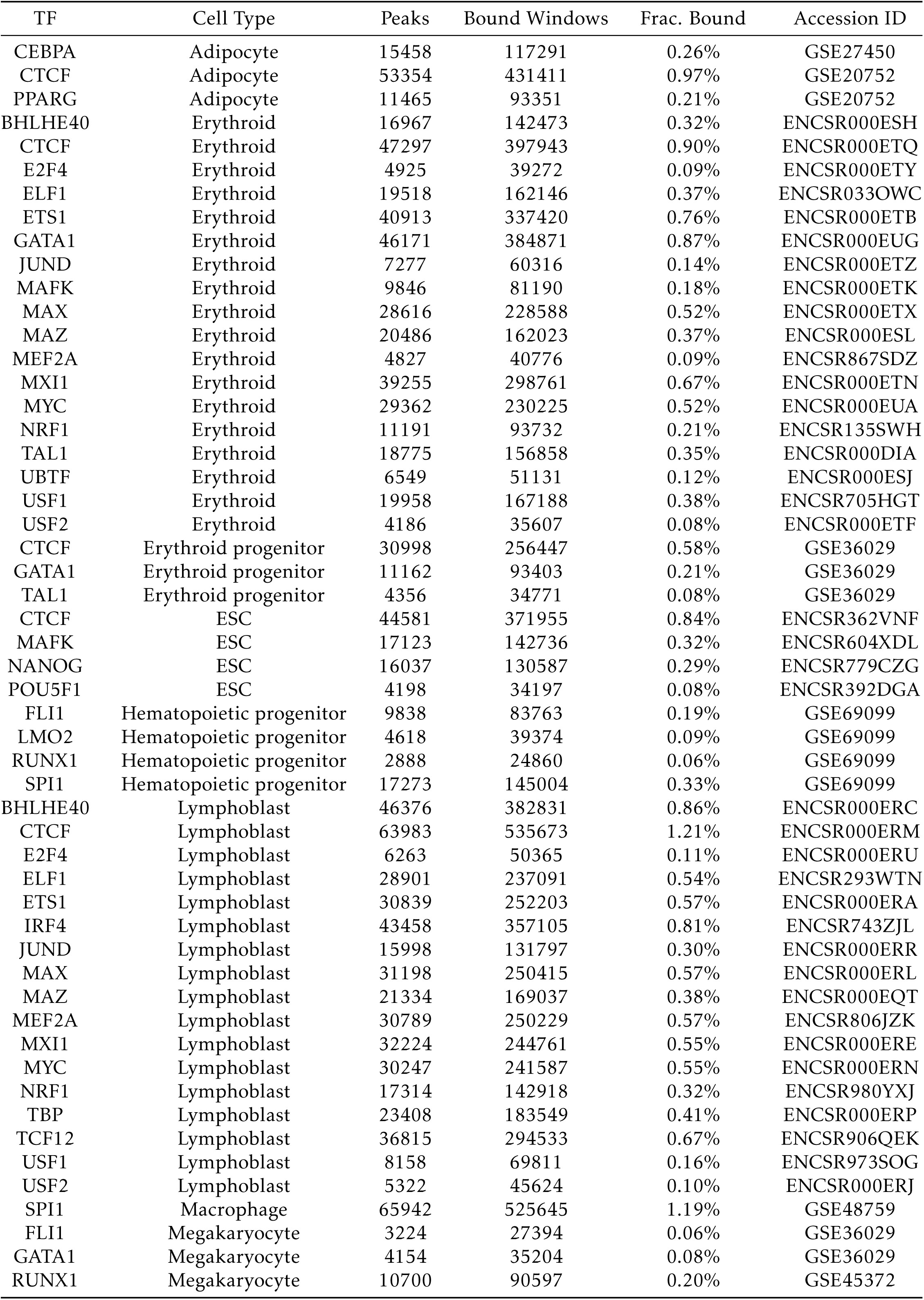
Summary statistics for all additional mouse datasets. The mouse genome filtered window set consisted of 44288170 windows in total.

**Table S7:** Summary statistics for all additional human datasets. The human genome filtered window set consisted of 51548966 windows in total.

| TF | Cell Type | Peaks | Bound Windows | Frac. Bound | Accession ID |
| --- | --- | --- | --- | --- | --- |
| CEBPA | Adipocyte | 53157 | 396024 | 0.77% | GSE27450 |
| CTCF | Adipocyte | 48914 | 376510 | 0.73% | GSE20752 |
| PPARG | Adipocyte | 58757 | 462122 | 0.90% | GSE20752 |
| BHLHE40 | Erythroid | 27808 | 217471 | 0.42% | ENCSR000EGV |
| CTCF | Erythroid | 59803 | 476076 | 0.92% | ENCSR000DMA |
| E2F4 | Erythroid | 9109 | 68965 | 0.13% | ENCSR000EWL |
| ELF1 | Erythroid | 32683 | 258940 | 0.50% | ENCSR000BMD |
| ETS1 | Erythroid | 13775 | 101997 | 0.20% | ENCSR000BKQ |
| GATA1 | Erythroid | 14676 | 113735 | 0.22% | ENCSR000EFT |
| JUND | Erythroid | 47180 | 367973 | 0.71% | ENCSR000EGN |
| MAFK | Erythroid | 27213 | 213251 | 0.41% | ENCSR000EGX |
| MAX | Erythroid | 37342 | 286474 | 0.56% | ENCSR000EFV |
| MAZ | Erythroid | 40398 | 308748 | 0.60% | ENCSR000EFX |
| MEF2A | Erythroid | 6407 | 49536 | 0.10% | ENCSR000BNV |
| MXI1 | Erythroid | 9081 | 70132 | 0.14% | ENCSR000EGZ |
| MYC | Erythroid | 31378 | 233216 | 0.45% | ENCSR000EGJ |
| NRF1 | Erythroid | 4436 | 36511 | 0.07% | ENCSR000EHH |
| TAL1 | Erythroid | 29476 | 229424 | 0.45% | ENCSR000EHB |
| UBTF | Erythroid | 19228 | 139064 | 0.27% | ENCSR000EFZ |
| USF1 | Erythroid | 22382 | 177524 | 0.34% | ENCSR000BKT |
| USF2 | Erythroid | 3621 | 29702 | 0.06% | ENCSR000EHG |
| CTCF | Erythroid progenitor | 36729 | 292844 | 0.57% | GSE26501 |
| GATA1 | Erythroid progenitor | 25710 | 198358 | 0.38% | GSE26501 |
| TAL1 | Erythroid progenitor | 38152 | 285562 | 0.55% | GSE26501 |
| CTCF | ESC | 57384 | 466110 | 0.90% | ENCSR000BNH |
| MAFK | ESC | 13422 | 109310 | 0.21% | ENCSR000EBS |
| NANOG | ESC | 8905 | 72332 | 0.14% | ENCSR000BMT |
| POU5F1 | ESC | 5029 | 41330 | 0.08% | ENCSR000BMU |
| FLI1 | Hematopoietic progenitor | 38760 | 310707 | 0.60% | GSE45144 |
| LMO2 | Hematopoietic progenitor | 2037 | 16312 | 0.03% | GSE45144 |
| RUNX1 | Hematopoietic progenitor | 29950 | 241749 | 0.47% | GSE45144 |
| SPI1 | Hematopoietic progenitor | 167273 | 1283083 | 2.49% | GSE70660 |
| BHLHE40 | Lymphoblast | 28651 | 227674 | 0.44% | ENCSR000DZJ |
| CTCF | Lymphoblast | 41765 | 339466 | 0.66% | ENCSR000DZN |
| E2F4 | Lymphoblast | 4375 | 35071 | 0.07% | ENCSR000DYY |
| ELF1 | Lymphoblast | 27369 | 212273 | 0.41% | ENCSR000BMB |
| ETS1 | Lymphoblast | 12912 | 103978 | 0.20% | ENCSR000BKA |
| IRF4 | Lymphoblast | 23043 | 182227 | 0.35% | ENCSR000BGY |
| JUND | Lymphoblast | 7602 | 61307 | 0.12% | ENCSR000DYS |
| MAX | Lymphoblast | 13605 | 104721 | 0.20% | ENCSR000DZF |
| MAZ | Lymphoblast | 23166 | 175906 | 0.34% | ENCSR000DZA |
| MEF2A | Lymphoblast | 22588 | 180702 | 0.35% | ENCSR000BKB |
| MXI1 | Lymphoblast | 21737 | 164076 | 0.32% | ENCSR000DZI |
| MYC | Lymphoblast | 4950 | 37375 | 0.07% | ENCSR000DKU |
| NRF1 | Lymphoblast | 3363 | 27933 | 0.05% | ENCSR000DZO |
| TBP | Lymphoblast | 19535 | 147978 | 0.29% | ENCSR000DZZ |
| TCF12 | Lymphoblast | 25023 | 201436 | 0.39% | ENCSR000BGZ |
| USF1 | Lymphoblast | 8461 | 69700 | 0.14% | ENCSR000BGI |
| USF2 | Lymphoblast | 4450 | 36621 | 0.07% | ENCSR000DZU |
| SPI1 | Macrophage | 88793 | 693731 | 1.35% | GSE31621 |
| FLI1 | Megakaryocyte | 4649 | 38182 | 0.07% | GSE24674 |
| GATA1 | Megakaryocyte | 4147 | 33052 | 0.06% | GSE24674 |
| RUNX1 | Megakaryocyte | 58757 | 209261 | 0.41% | GSE24674 |

